# Postnatal Reprogramming Shapes Human Intestinal Epithelial Immune Competency

**DOI:** 10.64898/2026.05.05.722861

**Authors:** Chloe Hyun-Jung Lee, David Fawkner-Corbett, Zoe Christoforidou, Ana Sousa Gerós, Verena Lentsch, Lucy Sheikh, Esther Bridges, Marta Jagielowicz, Lei Deng, Xiao Qin, Huei-Wen Chuang, Vy Wien Lai, Sophie Craddock, Aurore Mazurier, Paulina Siejka-Zielinska, Paula Gomez Castro, Anna Aulicino, Colleen McGregor, Tarun Gupta, Nicole Cianci, Rhea Kujawa, Paola Vargas Gutierrez, Carol Cheng, Maria Greco, Darren Fowler, Simon James Alexander Buczacki, Grace Rimmer, Rachel Harwood, Nigel Hall, Paul Johnson, Hashem Koohy, Alison Simmons, Agne Antanaviciute

**Affiliations:** Medical Research Council Translational Immune Discovery Unit (MRC TIDU), Weatherall Institute of Molecular Medicine (WIMM), University of Oxford, Oxford, UK; MRC WIMM Centre for Computational Biology, WIMM, Oxford, UK; Academic Paediatric Surgery Unit, Nuffield Department of Surgical Sciences, University of Oxford, Oxford, UK; Translational Gastroenterology and Liver Unit, John Radcliffe Hospital, Oxford, UK; MRC WIMM Advanced Single Cell OMICS Facility, WIMM, Oxford, UK; Paediatric Pathology, Department of Cellular Pathology, Oxford University Hospitals NHS Foundation Trust, Oxford OX3 9DU, UK; Nuffield Department of Surgical Sciences, Medical Sciences Division, University of Oxford; Alder Hey Children’s, Hospital, Liverpool, UK; School of Life Sciences, University of Liverpool, Liverpool, UK; University Surgery Unit, Faculty of Medicine, University of Southampton, Southampton, UK; Department of Paediatric Surgery and Urology, Southampton Children’s Hospital, Southampton, UK

## Abstract

At birth, the intestine must rapidly adapt to enable nutritional function and immune microbial tolerance. Here, integrating single-cell multi-omics and spatial transcriptomics we define the circuits underpinning this process. We identify asynchronous developmental trajectories with postnatal epithelial reprogramming characterised by coordinated changes in metabolism, junctional structure and innate defence. At birth epithelial stem cells demonstrate dynamic enhancer remodelling, with accessibility often preceding transcription. Fetal stemness elements remain accessible despite reduced transcription across epithelial lineages, retaining plasticity potential. Post-natal epithelia experience sequential homing of myeloid cells followed by innate T cells with peri-epithelial B cells localising later in infancy. Using developmentally staged organoids, we show that epithelial responses to inflammatory stimuli are age-dependent and constrained in early life. We identify *BHLHE40* as an early-life regulator that attenuates the impact of interferon- and NF-κB–driven signalling. Altogether we define the events driving epithelial licensing and barrier adaptation at birth and through infancy.

## Introduction

In the human intestine a myriad of specialized cell types and states act in concert to sustain digestion, absorption, immune regulation and barrier integrity. Coordination of these diverse cell states changes rapidly during the transition from fetal development to infant life, when the intestine becomes a primary mucosal interface to the external environment. At birth, the neonatal gut traverses through an abrupt environmental shift, encountering food antigens and undergoing colonisation by commensal microbes. Microbial colonization drives the rapid establishment of homeostatic molecular circuits that shape life-long barrier health ^1–3^. Defects in barrier maturation in postnatal life underpin a variety of conditions including necrotizing enterocolitis, neonatal sepsis and infectious diarrhoea^4–6^. More broadly, postnatal environmental exposures may impact on immune imprinting, setting intestinal immunoregulatory thresholds for the longer-term health^1,2,7–9^. Despite the importance of these processes for both immediate and future health, a high-resolution mechanistic understanding of early life intestinal development is lacking.

During early infancy, the intestine must function simultaneously as an effective barrier against microbial invasion and as a permissive interface that enables both intestinal immune training and establishment of tolerance^4^. In animal models this early postnatal period has been defined as a critical developmental window during which the gut exhibits regulated physiological permeability, enabling antigen sampling and microbial-driven immune education, with transient epithelial permeability supporting the induction of regulatory T cells, IgA production and innate lymphoid cell maturation in a period coined the “window of opportunity” ^10–12^. In humans, however, the timing, regional specificity and molecular features governing this transient permeability phase remain less clearly defined, in part due to the rarity of healthy tissue from this age range. Current multimodal single cell technologies enable capture of high-resolution data from such rare tissue to address the nature of epithelial reprogramming, immune colonisation, mesenchymal remodelling and the molecular events governing the host response to microbial colonisation.

Recent adoption of such approaches has begun to illuminate key aspects of human fetal intestinal development, including epithelial crypt-villus morphogenesis, maturation of neuromuscular circuits, and immune cell colonization with the formation of gut-associated lymphoid tissue (GALT)^13–16^. Complementary studies in fetal and adult tissues have mapped the regulatory landscapes that underpin development and mature intestinal functions using chromatin accessibility assays ^17,18^. However, the immediate postnatal window remains unexplored, as underscored by recent meta-analyses^19,20^ highlighting that very few neonatal samples have been profiled across the current body of single cell studies in the intestine. More recent work by de Sousa Casal *et al*. characterised duodenal development in infants aged six months and older^21^, but the immediate postnatal period remains poorly defined.

Here, we present a comprehensive single-cell and spatially resolved atlas of the developing human intestine, generated by whole-cell multi-omics to capture simultaneous scRNA, scATAC and CITEseq data from each cell. We further undertook subcellular resolution MERSCOPE spatial transcriptomics across fetal, neonatal and paediatric intestinal tissues. Leveraging this dataset, we constructed a multimodal molecular clock to trace early intestinal maturation. In the early postnatal period, we identified transient cell groups and molecular pathways that oversee epithelial barrier adaptation, mesenchymal remodelling and immune thresholds. We document post-natal stem cell imprinting and the regulatory networks guiding epithelial barrier maturation, changes in gut permeability, and the spatial niches influencing epithelial defence. We use age-stratified epithelial organoids to explore how post-natal epithelial imprinting affects epithelial reactivity to pro-inflammatory stimuli and identify a master-regulator of post-natal immune responses that downregulates epithelial inflammatory responses in the face of newly primed imprinted chromatin architecture. Altogether we present a data resource that will inform future studies of paediatric and adult intestinal disease pathogenesis.

### Spatiotemporal regulation of intestinal development in early life

To map spatiotemporal development of the human gut in early life, we profiled normal intestinal tissue from colon and ileum spanning developmental timepoints from fetal 9 post conceptual weeks (PCW), to neonates through to adulthood (**Figure 1A-B, Methods**). Using whole-cell multi-omics^22^, we captured mRNA expression and chromatin accessibility from the same cells across 54 samples, separately enriching for immune, epithelial and stromal compartments. In parallel, we profiled 23 tissue samples using MERSCOPE high-resolution *in situ* spatial transcriptomics (ST) using a custom 500-plex gut development panel, specifically designed using single-cell data to capture spatially and temporally resolved cell states across development (**Figure 1B**).

**Figure 1.**
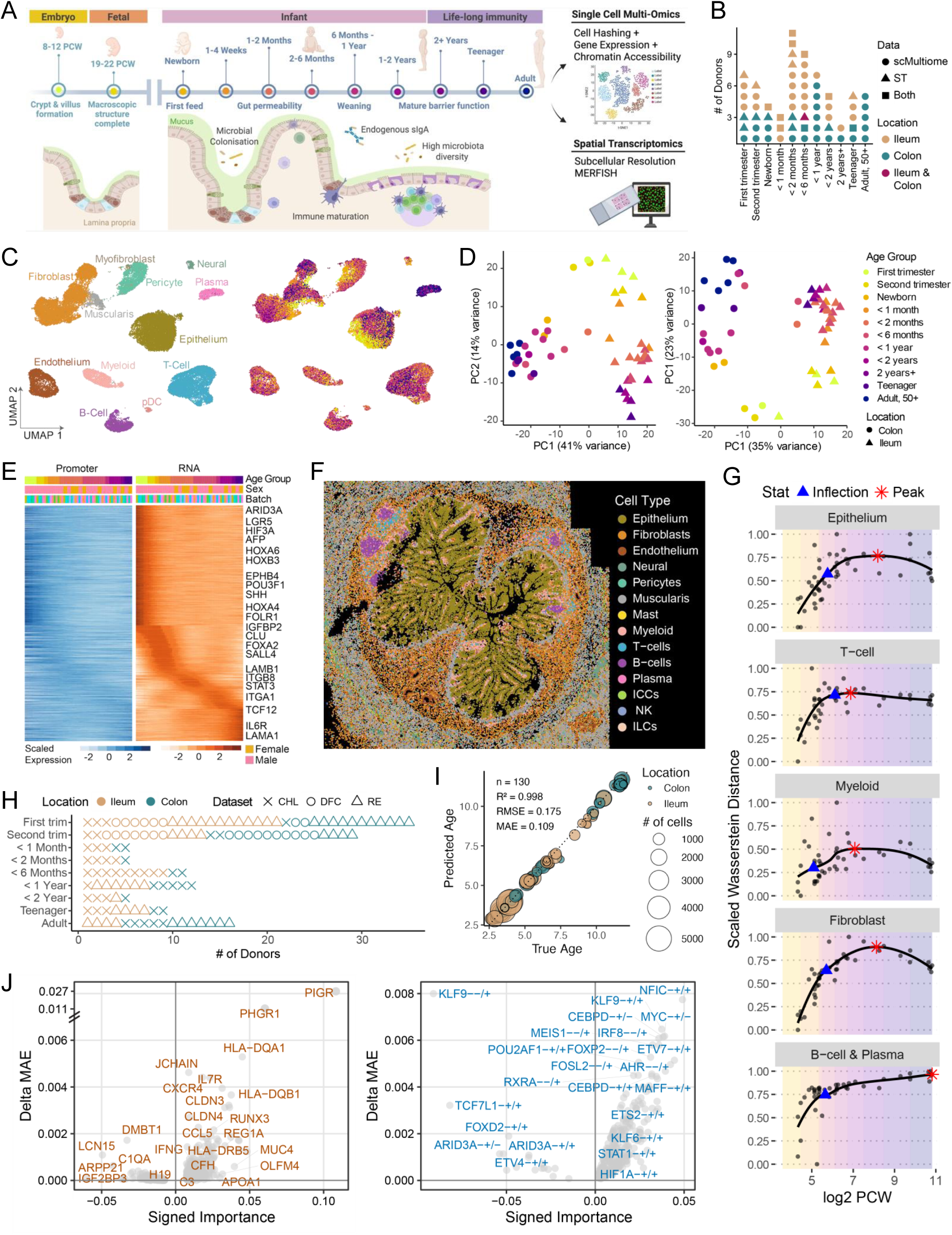
Spatiotemporal regulation of intestinal development in early life. A. Schematic of intestinal developmental timepoints analysed by single cell multiomics and in situ spatial transcriptomics (ST). Created with BioRender. B. Overview of single cell multiomics and ST cohorts; each point represents an individual donor. C. UMAP of integrated single cell multiomics data showing all cells, coloured by broad cell type and age group. Colour scheme consistent with D. D. Pseudobulk principal component analysis (PCA) of epithelial, stromal and immune compartments identified by single cell multiomics. Data were downsampled to equal cell numbers and coloured by age group. E. Heatmaps of promoter accessibility and gene expression for development-associated genes in ileal epithelium; values are mean-scaled per sample. Age group colour scheme consistent with D. F. Representative terminal ileum section (3 months) profiled by MERSCOPE ST, coloured by broad cell type. G. Scaled Wasserstein distance from a location-matched fetal anchor plotted against log2-transformed post-conceptual weeks (PCW). Each point represents an individual donor. LOESS fits are shown, highlighting the inflection and peak of the trajectories. H. Integrated transcriptomic atlas combining scRNA-seq data with datasets from Elmentaite et al. 2021 and Fawkner-Corbett et al. 2021; each point represents a donor. I. Performance of a multilayer perceptron (MLP) regressor on the validation dataset. Predicted age corresponds to the median across cells per donor. Points are coloured by location and sized by cell number. J. Volcano plots showing top genes (left) and regulons (right) from the colonic ElasticNet model. Axes indicate signed feature importance and change in mean absolute error (MAE) following in silico perturbation.

Following QC, we captured 235,465 cells using single cell multi-omics and 1,457,779 cells using *in situ* ST, covering all major epithelial, immune and stromal lineages (**Figure 1C-D**, **Supplementary Figure 1A-D**). Overall, we observed high agreement between the two technologies, with many cell populations exhibiting age-associated patterns (**Figure 1D, Supplementary Figure 1C, 2**). While epithelial populations displayed marked regional differences between the small and large intestine, non-epithelial lineages were comparatively conserved across locations (**Supplementary Figure 1C-D**).

To examine how transcriptional programmes are regulated across development, we compared mRNA expression with chromatin accessibility and observed strong concordance in both global structure and canonical marker profiles across major cell types (**Figure 1D-E, Supplementary Figure 1C, E-F**). Across development, genes showing age-dependent expression – including key developmental regulators such as HOX genes, components of SHH-WNT-BMP signalling pathways and lineage specifying transcription factors (TFs)^23^ – exhibited concordant changes in promoter accessibility, indicating that temporal transcription programmes are regulated through dynamic modulation of chromatin accessibility (**Figure 1E**). Our analyses further illustrated how coordinated promoter–enhancer regulation contributes to cell-state specification - for example, although the *F3* promoter was broadly accessible, a subset-specific enhancer was uniquely accessible in telocytes^24–26^, where *F3* is selectively expressed (**Supplementary Figure 1E-F**).

Given the complementarity of transcriptomic and epigenetic modalities, we defined cell states from joint RNA–ATAC embeddings, identifying 178 cell types and states distinguished by marker expression, chromatin accessibility profiles, differentiation status, developmental stage and anatomical location (**Supplementary Figure 2A, Supplementary Data**). To capture higher-order regulatory structure, we inferred TF regulatory networks and identified active regulons that recapitulated known lineage defining programmes (**Supplementary Figure 1G**).

The spatial data enabled us to resolve dynamic reorganisation of intestinal cellular architecture throughout maturation. We identified 139 distinct cell types and states in the spatial dataset, many of which exhibited pronounced age-associated differences consistent with the multi-omics analyses (**Supplementary Figure 2B**). Over developmental time, we observed a marked expansion in tissue size accompanied by increased architectural and cellular diversity and greater organisation complexity, reflecting progressive functional specialisation of the gut (**Figure 1F, Supplementary Figure 2C-D**).

### A molecular clock quantifies asynchronous intestinal cell maturation rates

Intestinal development involves sequential waves of cellular reprogramming which cannot be accurately represented by chronological age alone. To quantify developmental maturation at a molecular level, we measured how simultaneous scRNAseq and scATACseq profiles from neonatal and paediatric samples diverged relative to earliest fetal samples. This enabled measurement of intestinal maturation via an unbiased, data-driven distance metric (**Methods**). Both gene expression and chromatin accessibility showed a sharp divergence during late fetal and early neonatal stages, followed by a gradual stabilisation with age (**Supplementary Figures 3A-C**). These maturation trajectories peaked and plateaued at ∼2 years 9 months in the colon and ∼8 months in the ileum (**Supplementary Figure 3C-D**), indicating that the majority of age-associated changes occur within early postnatal life, with cellular reprogramming stabilising earlier in the ileum than in the colon.

To understand the basis of these regional differences, we examined which features were associated with developmental progression. Variation was driven in part by differences in cellular composition, with cell types such as B cells, which are more abundant in the colon, continuing to increase in prevalence and to undergo progressive transcriptional and phenotypic changes well beyond early life (**Supplementary Figure 3D, Supplementary Data**). This indicated that intestinal cell populations mature at different rates, leading us to examine maturation dynamics across individual cell types. Cell-type level analyses demonstrated substantial heterogeneity in developmental timing across intestinal cell lineages. Epithelial populations, together with myeloid and T-cell compartments, progressed through postnatal developmental states earlier than other cell types, whereas populations such as fibroblasts and B/plasma cells continued to undergo changes into later stages of life (**Figure 1G, Supplementary Figure 3E**). These findings indicate that while most intestinal cell types undergo coordinated development which largely stabilise post-infancy (**Supplementary Figure 2A**), colonic follicle biology remained dynamic into later childhood, reflective of ongoing establishment of adaptive mucosal immunity during this time.

Having established distance from a fetal reference as a quantitative measure of developmental state, we next asked whether this could be formalised into a predictive molecular clock model (**Supplementary Figure 3F-G, Methods**). Using single-cell multi-omics data from epithelial, stromal and immune compartments, we modelled developmental progression to allow comparison across donors and cell types, and to identify features most closely associated with intestinal maturation. We first trained a series of Elastic Net models using different combinations of molecular features derived from the scRNA and scATAC profiles. Among these, the model that incorporated TF regulon activity in addition to gene expression was most effective at predicting intestinal maturation state (Mean absolute error (MAE)= 0.438 for colon and 0.728 for ileum) (**Supplementary Figure 3F-G**).

Although the multimodal model performed well within our cohort, representation of early fetal stages was limited. To extend developmental coverage and improve resolution around birth, we integrated published sc RNA-seq datasets spanning fetal to adult stages. Using this expanded transcriptomic atlas, we trained a secondary multilayer perceptron (MLP) regression model (MAE=109), which is well suited to capturing non-linear developmental trends. This analysis validated and refined maturation trajectories across donors and reinforced the concentration of major molecular transitions within early postnatal life (**Figure 1H-I, Supplementary Figure 4A-B**).

In both models, features predictive of later developmental time were enriched for immune-associated programmes, including immunoglobulin expression and antigen-presentation markers, consistent with progressive immune maturation. In contrast, earlier developmental states were characterised by fetal-associated stromal and epithelial features and by TF regulatory programmes enriched during early life. Within the epithelial compartment, the molecular clock model highlighted early-life regulons such as ARID3A, alongside transient expression of fetal markers including *TTR* and *AFP* (**Figure 1J, Supplementary Figure 4C-D**).

Together, these analyses identify early neonatal life as a period of pronounced regulatory change and highlight epithelial-associated programs as key drivers of intestinal remodelling during infancy.

### Functional reprogramming of the epithelium at birth and over infancy

To explore infant epithelial adaptations, we first examined how epithelial function shifts across development in response to changing physiological demands. Examining scRNAseq data demonstrated fetal epithelium was enriched for metabolic pathways supporting cellular growth and biosynthesis, including amino acid and vitamin metabolism, consistent with rapid intrauterine expansion (**Figure 2A, Supplementary Figure 5A-B**). After birth, pathway activity switched towards lipid utilisation, redox regulation and nutrient absorption, reflecting the transition to enteral feeding and microbial exposure. Lipid processing programmes activated early in postnatal life included a transient increase in *CEL* expression, which facilitates digestion of breast milk-derived fats. Conversely, carbohydrate metabolism pathway genes, such as *LCT*, *MGAM* and *SI*, increased progressively during infancy (**Figure 2A, Supplementary Figure 5A-C**), coinciding with weaning and dietary diversification.

**Figure 2.**
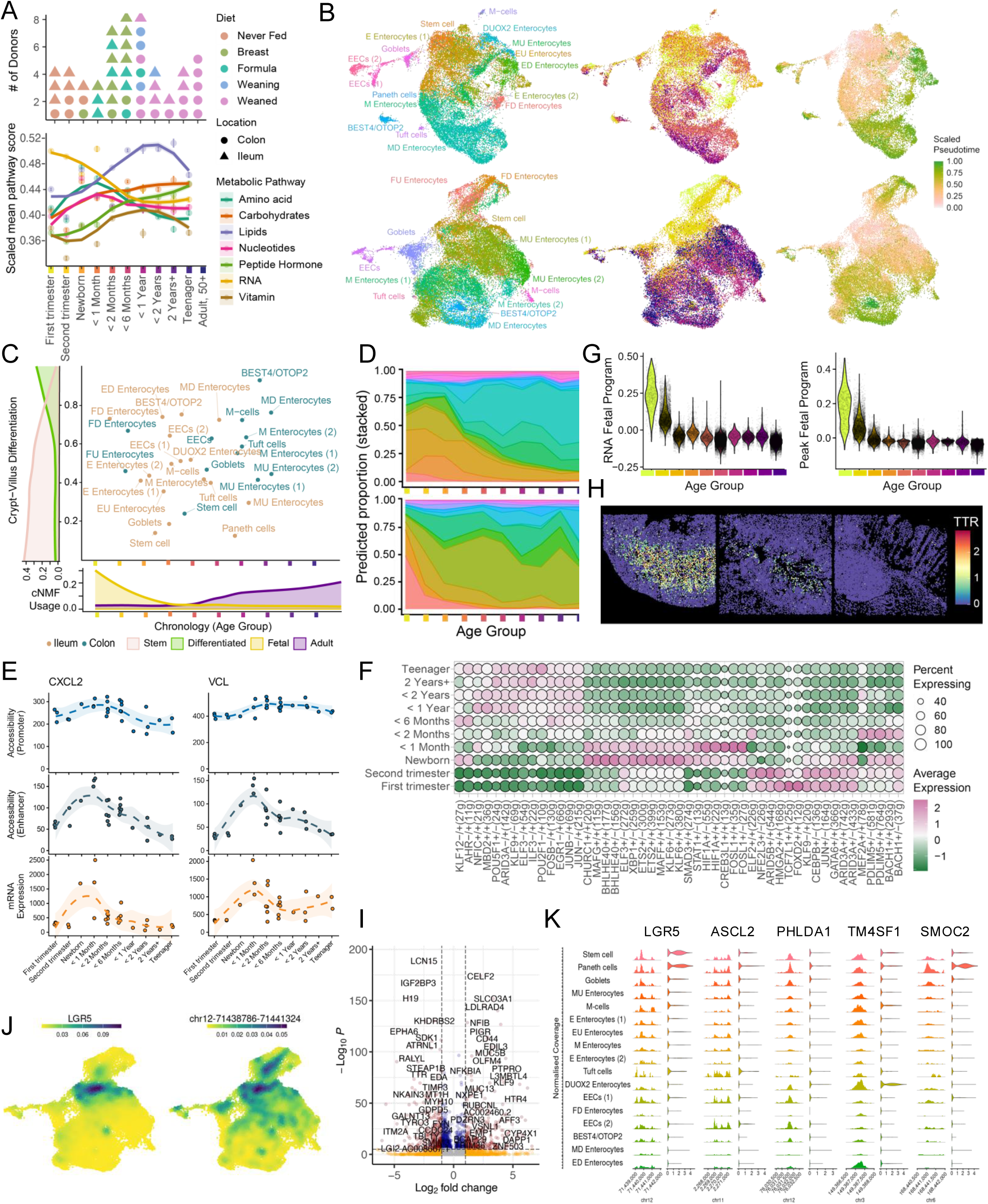
Diet, differentiation and chromatin dynamics shape epithelial maturation. A. Top, overview of diet conditions in single cell multiomics cohorts; each point represents a donor, coloured by diet and shaped by location. Bottom, scatterplot with LOESS-fitted line showing metabolic pathway scores in ileal samples, coloured by pathway. B. UMAP of ileal (top) and colonic (bottom) epithelial cells in a joint RNA–ATAC embedding, coloured by cell type, age group (as in A) and crypt–villus differentiation pseudotime. C. Relationship between mean chronological age (by age group) and crypt–villus differentiation score across epithelial cell clusters, coloured by location. Distributions show stem, differentiated, fetal and adult cNMF module usage across chronology (age group) and differentiation. D. Predicted proportions of epithelial cell types in ileum (top) and colon (bottom), estimated using sccomp and visualized as ribbon plots; colours correspond to cell types in B. E. Generalized additive model (GAM)-fitted trends of promoter and enhancer accessibility per sample and gene expression for selected genes across age in ileal epithelium. Shaded areas indicate +/- 1 standard error of the GAM fitted mean. F. Bubble plot showing age-associated regulon activity across ileal epithelial cells. G. Violin plots showing fetal gene module scores (left) and chromatin accessibility module scores (right) in colonic epithelium. H. Spatial expression of the fetal marker TTR in MERSCOPE ST sections at 9 PCW (left), 5 days (middle) and 13 months (right). I. Volcano plot showing differential expression between fetal and paediatric colonic stem cells in single cell multiomics data. Differential expression was assessed using DESeq2 (Wald test with Benjamini–Hochberg correction). Points are coloured by significance and effect size (red, |log2FC| > 1 and −log10P > 1; blue, |log2FC| < 1 and −log10P > 1; yellow, |log2FC| > 1 and −log10P < 1; grey, not significant). J. Density plots of LGR5 gene expression (left) and promoter accessibility (right) in colonic epithelial cells. K. Coverage and violin plots showing promoter accessibility and gene expression of stem cell marker genes in ileal epithelium, coloured by mean crypt–villus differentiation score per cell type.

These environmentally driven neonatal and infant metabolic adaptations were also reflected in epithelial cell-state composition across development. We identified stem, secretory and absorptive epithelial cell populations at all ages sampled (**Figure 2B**). Across these lineages, age explained the largest proportion of variance in epithelial cell states, exceeding associations with intestinal location or diet (**Supplementary Figure 5D**). Several epithelial clusters were enriched at specific developmental stages, including fetal-associated enterocyte populations and early neonatal clusters that were more prevalent in neonates and young infants (**Figure 2C-D**). However, these clusters did not segregate strictly by age, instead showing gradual tapering across developmental windows; for example, late fetal enterocyte cell states were still detectable in early neonatal samples. These patterns were also reflected in the activity of gene expression programs detected by cNMF analysis^27^ (**Supplementary Figure 5E-H**), where we identified gene programmes associated with fetal and adult epithelial states, as well as location and diet liked programs.

We observed marked separation of fetal and early life epithelial cell clusters (**Figure 2B**) which prompted us to examine whether these developmentally restricted epithelial states represent an increased proportion of undifferentiated intermediates or functionally distinct, developmental-window restricted differentiated cell states. To disentangle differentiation state from developmental time, we modelled epithelial cells along two orthogonal axes: crypt-villus differentiation and chronological developmental progression (**Methods**). Differentiation trajectories were computed from stem to mature absorptive and secretory states across all ages; however, fetal and early postnatal enterocytes remained transcriptionally distinct even after accounting for differentiation state. Accordingly, stemness vs differentiation and fetal vs adult cNMF programmes captured two orthogonal axes of epithelial cells (**Figure 2C, Supplementary Figure 5E-H**). Thus, differentiated epithelial cells are present from early fetal life, but their functional programmes vary with developmental age. This indicates early life epithelium is not more “stem-like” but rather reflects age-specific functional reprogramming aligned with changing physiological and environmental demands.

### Regulatory architecture of early life epithelial cell state transitions

We next examined how early life epithelial state transitions are encoded at the chromatin level. We first asked whether chromatin accessibility corresponds to transcriptional output by calculating Wasserstein distance (**Methods, Supplementary Figure 6A**) between RNA and ATAC embeddings of the same cells, highlighting that these modalities do not always agree and capture complementary aspects of cell state. Across epithelial cells, we observed multiple modes of relationship between chromatin accessibility and gene expression. In many cases, RNA and ATAC were concordant: for example, accessibility at the *ITGA2* promoter and enhancer closely tracked transcriptional peak in early postnatal samples; genes associated with weaning period, such as *SLC30A10* also showed similar concordance (**Supplementary Figure 6B**). However, we also identified genes in which promoter accessibility remained constitutively open across development while transcriptional dynamics were driven by distal regulatory elements. For instance, *VCL* and *CXCL2* showed stable promoter accessibility from fetal stages, whereas postnatal increases in expression were associated with increased enhancer accessibility (**Figure 2E**). Similarly, *GALNT2* expression declined with reduced enhancer activity despite unchanged promoter accessibility (**Supplementary Figure 6B)**. Thus, temporal regulation is frequently mediated by enhancer dynamics rather than promoter accessibility alone; indeed, enhancer remodelling has previously been demonstrated to drive epithelial stem cell locational identity^28^.

We further identified genes in which chromatin accessibility changed independently of transcription. For example, the promoter of *CXCL9*, an interferon-inducible gene with very limited expression in our healthy sample cohort, was largely inaccessible in fetal epithelium but became progressively accessible after birth (**Supplementary Figure 6B)**. This suggests that chromatin accessibility encodes transcriptional “readiness”, such that fetal and postnatal epithelial cells may differ in their capacity to induce, for example, pro-inflammatory responses upon stimulation.

We next examined TF regulatory networks that integrate accessible chromatin with TFs and their downstream target gene expression (**Methods**). We identified regulons associated with differentiation, metabolism and immune function, several showing strong age dependence (**Figure 2F**). Early-life epithelial cells were characterised by increased activity of developmental regulators such as ARID3A, whose targets are enriched for epithelial growth and proliferation functions^29^. *In silico* perturbation of these early-life regulons substantially impaired performance of our molecular age model, indicating that these programmes are important regulators of early epithelial cell identity (**Supplementary Figure 6C**).

Notably, several such fetal-associated transcriptional programmes and regulons persisted transiently after birth (**Figure 2A, 2F-G, Supplementary Figure 5A-B, Supplementary Figure 4D, Supplementary Figure 6D**). Expression of fetal genes such as *TTR*, involved in the transport of vitamin A, declined rapidly post-birth but nonetheless remained detectable in early neonatal samples. Spatial analysis revealed that, where such expression persisted, it became restricted to the crypt top and villus (**Figure 2H**). This pattern suggests that fetal gene expression post-birth may be restricted to differentiated progeny only and not maintained in progeny of postnatally imprinted stem cells.

To investigate how stem cells are reprogrammed at birth, we next examined regulatory differences within stem cell compartment over developmental time. Previous work in mice has shown that embryonic and adult intestinal stem cells have distinct molecular profiles^30^. Here, clustering analysis also segregated stem cells into fetal and postnatal groups, with postnatal stem cells upregulating metabolic and immune and defence programmes (*PIGR*, *NFKB1*, *NFKBIA*, *IFI6*), while fetal stem cells were enriched in developmental pathways (*TTR*, *H19*, *CLU*, *IGF2BP3*) (**Figure 2I, Supplementary Figure 6E-F, 7A-C**). Fetal stem cells failed to express *OLFM4* which was observed only after birth^31^. Together, this suggests rapid postnatal reprogramming of stem cells during the window of initial microbial exposure, with these defence programmes subsequently propagated to differentiated daughter cells.

Notably, separation between developmental stages was more pronounced in chromatin accessibility than in transcriptional space, with stem cells from older donors exhibiting greater divergence in ATAC profiles than RNA profiles (**Supplementary Figure 6E**). At individual loci, accessibility of key stem and regeneration-associated genes was maintained independently of expression. For example, *CLU*, previously implicated in revival stem cell states ^32–34^, remained accessible across developmental stages despite declining expression after birth (**Supplementary Figure 7D**). This suggests that fetal-associated stem cell programmes are not fully erased during postnatal maturation and that this retention in accessibility provides a potential epigenetic mechanism for reactivation of fetal-like regenerative programmes during injury.

More broadly, we found that accessibility of stem cell–associated regulatory elements extended well beyond the stem cell compartment. Chromatin at stemness-associated loci (*LGR5*, *ASCL2*, *PHLDA1*, *TM4SF1*, *SMOC2*) remained accessible across multiple differentiated epithelial lineages despite minimal gene expression outside of stem cells and was only substantially reduced in terminally differentiated enterocytes (**Figure 2J-K, Supplementary Figure 7E-G**). This discordance between accessibility and transcription indicates that the majority of epithelial cells retain a latent regulatory competence to activate stem-associated programmes; this is in line with observations that many epithelial cells are capable of de-differentiation during injury ^33,35–41^, and our data suggests that only terminally differentiated enterocytes lose this regenerative plasticity. Notably, we also observed this pattern in differentiated fetal enterocytes.

### Postnatal regulation of epithelial barrier permeability

The first few weeks of life is a period of profound environmental changes within the neonatal intestine, when the epithelium encounters nutrients, food antigens and microbial communities for the first time. In animal models this early postnatal period has been defined as a critical developmental window during which the gut exhibits regulated physiological permeability, enabling antigen sampling and microbial-driven immune education, with transient epithelial permeability supporting the induction of regulatory T cells, IgA production and innate lymphoid cell maturation^12^. In humans, however, the timing, regional specificity and molecular regulation of this “window of opportunity” permeability phase remain far less clearly defined.

Tight junctions control this process through coordinated expression of cell adhesion molecules, scaffold proteins and polarity regulators; thus, we examined the expression of leaky and sealing claudins (members of the claudin family that respectively increase paracellular ion permeability or reinforce barrier tightness), as well as additional tight junction–associated genes (**Figure 3A, Supplementary Figure 8A-B**)^42^. Several genes (e.g. polarity regulator *SHROOM2*) were highly expressed during fetal development, peaked neonatally and declined within the first postnatal month. The leaky claudin *CDLN2* exhibited this expression pattern in colon, but not ileum – it was rapidly suppressed within one month postnatally in colon, but expression persisted for several months in ileum. *NECTIN2* was upregulated at birth in ileum but not colon. Scaffold proteins (*TJP1*, *OCLN*) and integrins also exhibited dynamic trajectories; *ITGB4* was upregulated at birth in both colon and ileum, while conversely, *TJP1* and *OCLN* were more highly expressed in fetal development. Sealing claudins *CLDN3/4* were constitutively expressed throughout the developmental time lime and were not downregulated at birth.

**Figure 3.**
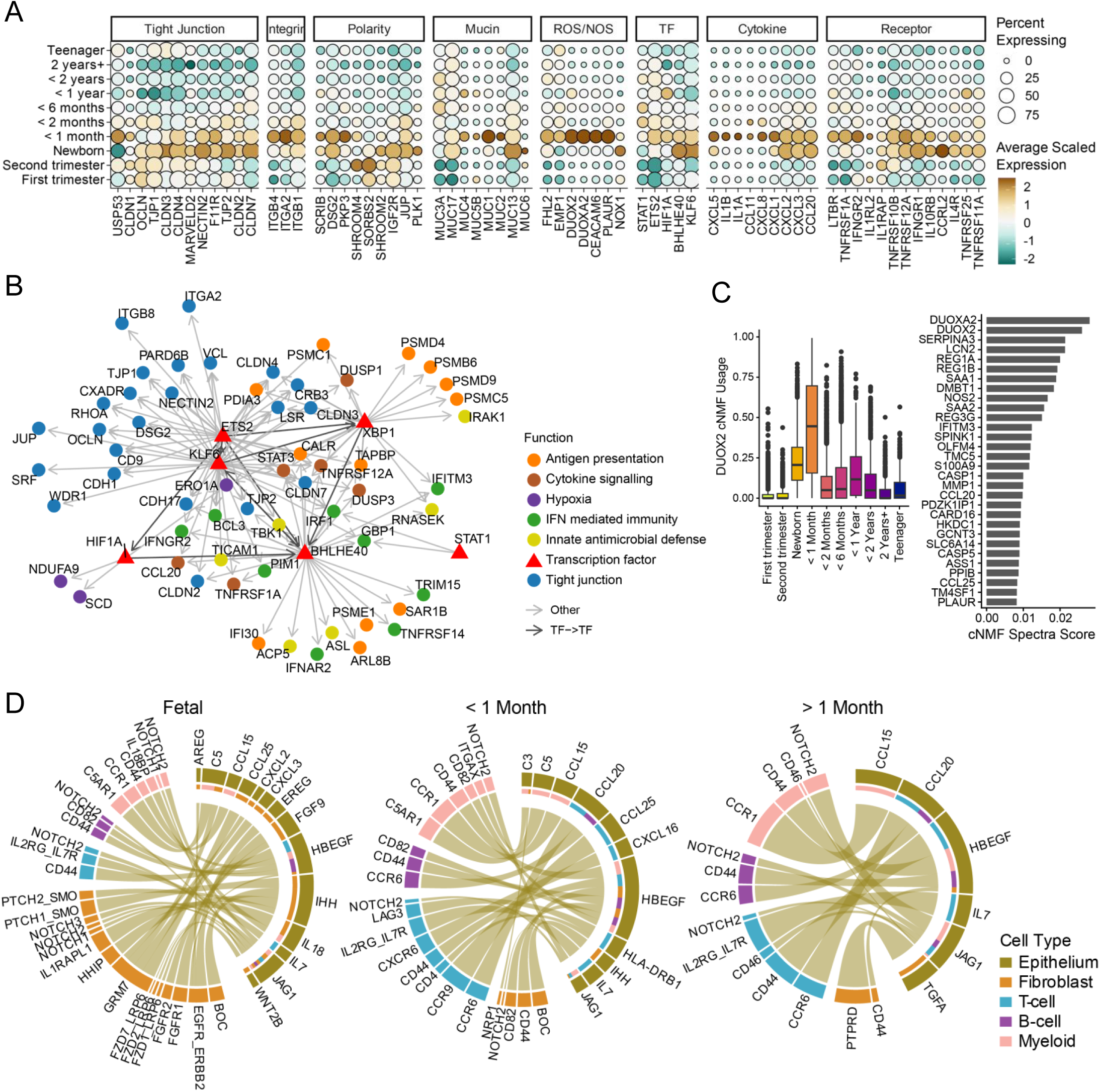
Activation of epithelial-intrinsic innate defence programmes in early life. A. Bubble plot showing expression of genes associated with tight junctions, integrins, polarity, mucins, reactive oxygen species (ROS), nitric oxide (NO), transcription factors, cytokines and receptors in ileal epithelial cells. B. Gene regulatory network showing transcription factors and their downstream targets (arrowed), coloured by functional pathway. C. Distribution of DUOX2-associated cNMF programme usage across age groups. Bar plot shows spectra score of top DUOX2 cNMF marker genes. Box plots indicate quartiles (Q1, median, Q3). D. Chord diagrams showing epithelial-to-immune and stromal cell communication inferred from ligand–receptor interactions. Interactions were subset by age group, ranked by log fold change (LIANA), and the top 40 interactions per group are shown. Chord width reflects interaction strength and colours denote cell types. Selected pathways include CCL, complement, CXCL, EGF, FGF, Hedgehog, IL1, IL2, MHC-II, NOTCH and WNT.

We next asked how these processes are regulated. TF regulatory network analysis identified two key regulators of junctional assembly and cell adhesion genes – *KLF6* and *ETS2*. *KLF6* and *ETS2* TF networks exhibited both high TF expression and downstream target activity during the first postnatal month (**Figure 2F, 3A-B, Supplementary Figure 8B**). Their downstream targets, many of which are enriched during this period, are involved in junctional assembly and cell adhesions (**Supplementary Figure 8C, Supplementary Data).** Together, this suggests that early postnatal barrier is not defined by a single, uniform phase of permeability, but by multiple time and region-specific programmes modulating junction components likely driven by *ETS2* and *KLF6*.

### Activation of epithelial-intrinsic innate defence programmes in early life

In parallel with the epithelial junction changes, we also observed upregulation of innate defence programs in early postnatal epithelium, in line with first exposure of epithelial cells to microbial stimuli^43–45^. We observed increased expression of *DUOX2*, *DUOXA2* and *NOX1* during the first month of life in ileum, along with an enrichment of DUOX2⁺ enterocyte cluster characterized by host defence genes (*DUOX2*, *DUOXA2*, *CEACAM6*, *C4BPB*) and tissue remodelling genes (*PLAUR*, *FHL2*, *EMP1*) (**Figure 3A, 3C, Supplementary Data**). This response likely contributes to mucosal defence through DUOX2-derived hydrogen peroxide and other antimicrobial ROS that help limit microbial overgrowth; commensal bacteria can also induce epithelial DUOX2-mediated ROS production to support barrier maturation^46^. Alongside *DUOX2* upregulation, the neonatal epithelium displayed increased expression of genes involved in redox regulation and cellular responses to oxidative stress **(Figure 3A, Supplementary Figure 8B)**. Notably, in our data, this period was characterised by an overlap in antioxidant systems, with relative enrichment of thioredoxin-associated components (*PRDX1/2*, *TXNRD1*) in fetal and early neonatal stages and a peak of glutathione-related pathways (*GPX2*) at birth (**Supplementary Figure 8D**). Glutathione pathway primarily detoxifies peroxides, while thioredoxin participates in redox regulation of TFs and signalling and is essential for mammalian embryonic development^47^. Thus, during the early neonatal period, epithelial redox pathways balance antimicrobial ROS production with antioxidant homeostasis and thioredoxin-dependent modulation of redox-sensitive TFs ^48^.

Concomitant with ROS and NOS pathway activation, expression of several mucins was also upregulated at birth. These included transmembrane mucins (*MUC1*, *MUC3A*, and *MUC17*) in both colon and ileum (**Figure 3A, Supplementary Figure 8A-B)**, which are membrane-anchored and not only restrict microbial access but can participate in immunomodulatory signalling^49^. *MUC1* can both attenuate excessive TLR signalling and activate pro-inflammatory signalling by increasing NF-kB activation and increasing production of pro-inflammatory cytokines, such as IL8 (*CXCL8*) ^50,51^.

In line with this, our regulon analysis further identified TFs associated with antimicrobial defence (*XBP1*), cytokine signalling (*STAT1*) and oxidative stress responses (*HIF1A*) activated in neonatal epithelium, together with corresponding elevated expression of their downstream target pro-inflammatory cytokines and chemokines including *CXCL1/2/3/5/8*, *CCL20* and *IL1B* (**Figure 3A, Supplementary Figure 8B**)^52^. This cytokine profile is dominated by molecules that preferentially recruit and activate myeloid cells and other innate immune cells (**Figure 3D, Supplementary Data**), as opposed to adaptive immune cells during this window.

### Evolving crypt microenvironment shapes postnatal epithelial responses

Epithelial induction of defence programmes during neonatal life occurs both within the context of microbial colonisation and a dynamically evolving crypt microenvironment. Therefore, we next asked how epithelial responses co-develop with the local stromal and immune signalling landscape. Within the mucosa (as identified by niche analysis, **Methods**), we observed a progressive transition from sparse immune representation during fetal development to an immune-enriched landscape postnatally, with a strong shift from innate to adaptive immunity through infancy (**Figure 4A-B, Supplementary Figure 9A-C**). Myeloid cells dominated the fetal mucosa and the first post-natal week (**Figure 4A-C**); T-cells accumulated in the mucosa after the first week, but did not diversify until 1 month after, while B-cells were largely absent during the first month and steadily accumulated throughout later infancy and early childhood (**Supplementary Figure 9A-C, Supplementary Figure 10A**).

**Figure 4.**
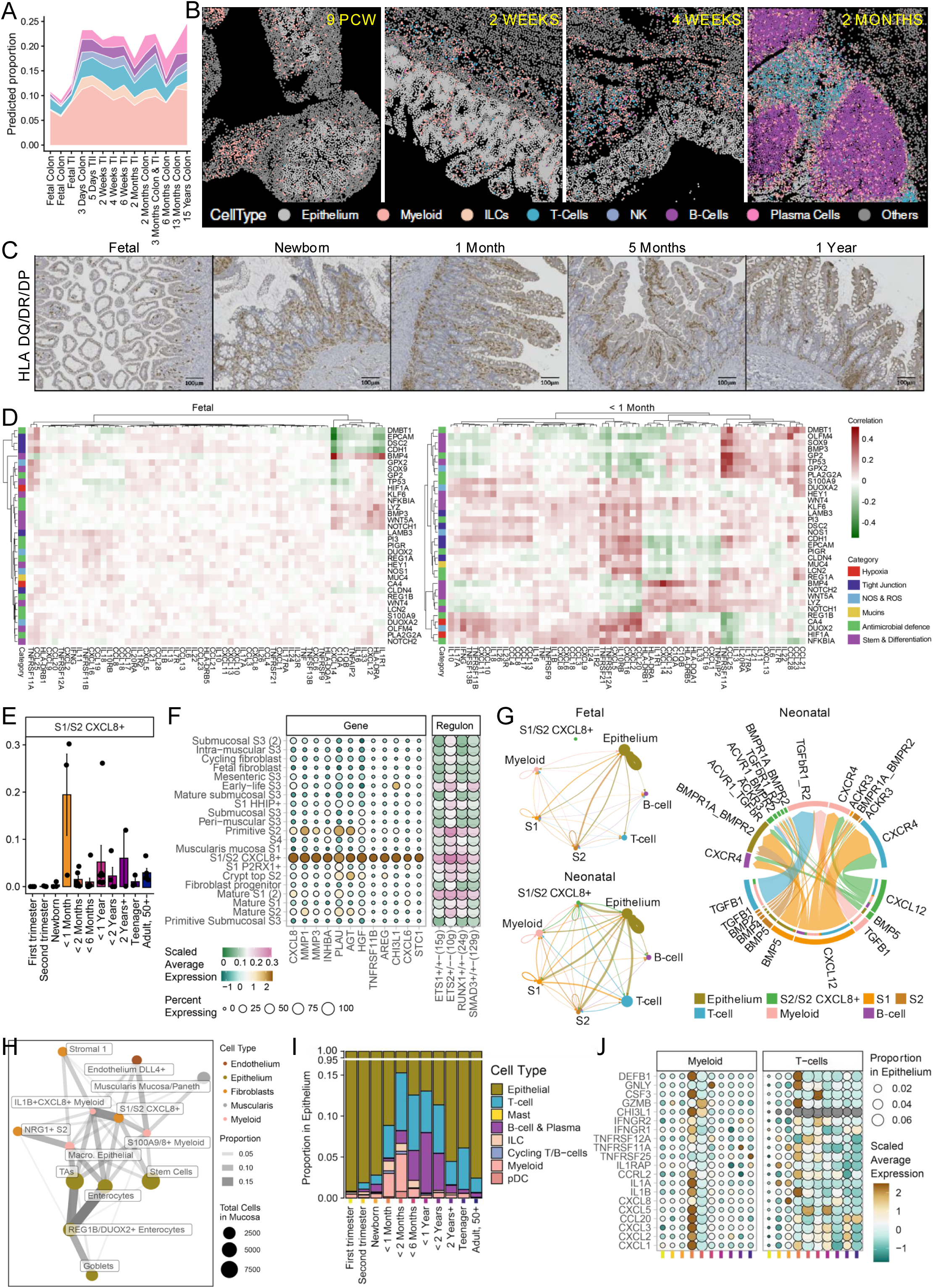
Evolving crypt microenvironment shapes postnatal epithelial responses. A. Predicted proportions of immune cell types across age groups in MERSCOPE ST data, estimated using sccomp and visualized as ribbon plots (myeloid, ILC, T cells, NK cells, B cells and plasma cells). Colours correspond to cell types in B. B. Representative MERSCOPE ST sections at 9 PCW, 2 weeks, 4 weeks and 2 months, coloured by cell type. C. Representative immunohistochemistry (IHC) images across developmental stages (fetal, newborn, 1 month, 5 months and 1 year) stained for HLA-DR. Scale bars, 100 μm. D. Heatmaps of epithelial–neighbourhood correlations across developmental stages. Pearson correlations between epithelial gene expression programmes and neighbouring cytokine-related signals were computed in terminal ileum samples, averaged within age groups (fetal, <1 month), and visualized by hierarchical clustering using a consistent colour scale. E. Proportion of S1/S2 CXCL8⁺ fibroblasts across age groups in single cell multiomics data. F. Bubble plot showing marker gene expression and regulon activity in S1/S2 CXCL8⁺ fibroblasts in single cell multiomics data. G. Cell–cell communication networks inferred using CellChat from single cell multiomics data. Left, circle plot showing interaction strength between epithelial, immune and fibroblast populations in fetal and neonatal (<2 months) samples. Right, chord diagram highlighting ligand–receptor interactions in CXCL, TGFβ and BMP signalling pathways. H. Cluster–neighbour co-localisation network from MERSCOPE ST data highlighting interactions among S1/S2 CXCL8⁺ fibroblasts, IL1B⁺CXCL8⁺ myeloid cells, REG1B/DUOX2⁺ enterocytes, S100A8/9⁺ myeloid cells and epithelial macrophages. Edges represent co-localisation strength (threshold > 0.02). Node size reflects cell abundance and colour denotes cell type. Data are from terminal ileum samples aged <1 month. I. Proportion of intraepithelial immune cells in chelated epithelial crypt single cell multiomics data across age groups. J. Bubble plot showing cytokine and immune state marker gene expression in intraepithelial immune cells; point size indicates the proportion of each cell type within age groups.

We next asked whether the unique epithelial innate defence programs we observed in early life coincided with the changing immune microenvironment by examining spatial gene expression correlations between epithelial programmes and cytokines and chemokines (**Figure 4D, Supplementary Figure 10B**). In samples from birth through the first month, we observed strong spatial co-expression between pro-inflammatory and alarmin-associated cytokines (*IL1B*, *IL33*), chemokines (*CXCL1/2*, *CXCL5/6, CXCL8, CCL20*), regulatory cytokines (*IL24*), and epithelial stress programmes marked by HIF1A and NFκB activity. These regions also showed co-localisation with *DUOX2* and antimicrobial defence signatures. *IFNG* expression spatially aligned with NOS/ROS programmes, consistent with local interferon signalling at the barrier. *KLF6*, a regulator of tight junction genes, strongly co-localised with these cytokine-rich zones in early life. These spatial couplings were largely absent in fetal tissue, diminished after the first postnatal month and substantially reduced beyond six months of age.

Examining the source of local crypt niche cytokine signalling in spatial data, we identified myeloid cells, T-cells and fibroblasts as a major cell type source of these signals (**Supplementary Figure 10C**). Multi-ome data confirmed a peak in pro-inflammatory cytokine signalling at birth and within the first month of life. At birth, M1-polarised macrophages increased and were the source of *IL1B*, *TNF* and *CXCL8*, while T-cell derived cytokines, including *IFNG*, peaked later after the first postnatal week (**Supplementary Figure 10C-D**). We further identified a neonatal-restricted mucosal fibroblast population enriched in samples < 1month of age (**Figure 4E-G).** These cells co-expressed both pro-inflammatory chemokines (*CXCL1*, *CXCL6*, *CXCL8*) and regulatory cytokines (*IL24*). Ligand–receptor analysis predicted active communication between these fibroblasts and immune and epithelial compartments during the neonatal stage, including CXCL12–CXCR4, CXCR1-CXCL8 and TGFB1–TGFβR interactions with myeloid and T cells and BMP5–BMPR signalling with epithelial cells (**Figure 4G-H**). This suggests that neonatal fibroblasts likely contribute to recruitment and positioning of myeloid cells in proximity to the epithelium during the first weeks of life.

In the multiome data, cells isolated from chelated epithelial crypt fractions enriched not only for epithelial cells but also for intraepithelial immune cells, providing a unique opportunity to query epithelial-associated immune populations in detail. Analysis of these cells revealed age-dependent changes in epithelium-associated immune composition (**Figure 4I**). Immune cells accumulated within crypts during the first postnatal month, with enrichment of myeloid cells, ILCs and T cells over the first two months, followed by expansion of B and plasma cells and a relative decline in myeloid populations over the first two years of life. Crypt-associated immune cells in early life displayed elevated expression of pro-inflammatory cytokines including *IL1A*, *IL1B*, *CCL20* and *CXCL2*, alongside activated inflammatory myeloid cells and GNLY⁺/GZMB⁺ cytotoxic T cells (**Figure 4I-J, Supplementary Figure 10C**). While this was largely consistent with changes in whole tissue CD45+ cells, comparison of gene activity programs between epithelial and total immune fraction T-cells revealed compartmental biases (**Supplementary Figure 10E**). γδ, TRM (Tissue-resident memory) and TEMRA (Terminally Differentiated Effector Memory T-cells) signatures were enriched among intraepithelial lymphocytes (IELs) across ages and increased over developmental time, while Th17 programmes were more prominent in lamina propria T cells. Naïve programs dominated early samples in both compartments, whereas cytotoxic and interferon-response programmes peaked within the first month (but not at birth) in both IELs and lamina propria T cells, with slightly stronger interferon signatures in IELs, suggesting enhanced local IFNγ signalling at the epithelial interface.

Together, these data show transiently elevated niche pro-inflammatory signalling within the first month of life, initially dominated by activated myeloid populations and subsequently shaped by a diversifying T-cell compartment and activated crypt niche fibroblasts. These observations are in line with previous studies that described heightened pro-inflammatory states in immune cells in infants ^53^.

### Organoids model age-dependent immune capacity of epithelial cells

Given the spatially restricted epithelial-immune dynamics observed in early life, together with marked age-dependent transitions in epithelial stem cell chromatin accessibility, we explored whether age dependent reprogramming impacted epithelial ability to respond to pro-inflammatory stimuli. To this end, we established human primary intestinal epithelium derived ileal organoids (PDOs)^54^ from fetal, neonatal and pediatric samples spanning six time points (**Methods, Figure 5A**). Spheroids were differentiated into budding organoids and subsequently stimulated with pro-inflammatory cytokines (**Figure 5B, Methods**). IFNγ or TNF and bulk mRNA sequencing was performed to profile age-dependent transcriptional responses.

**Figure 5.**
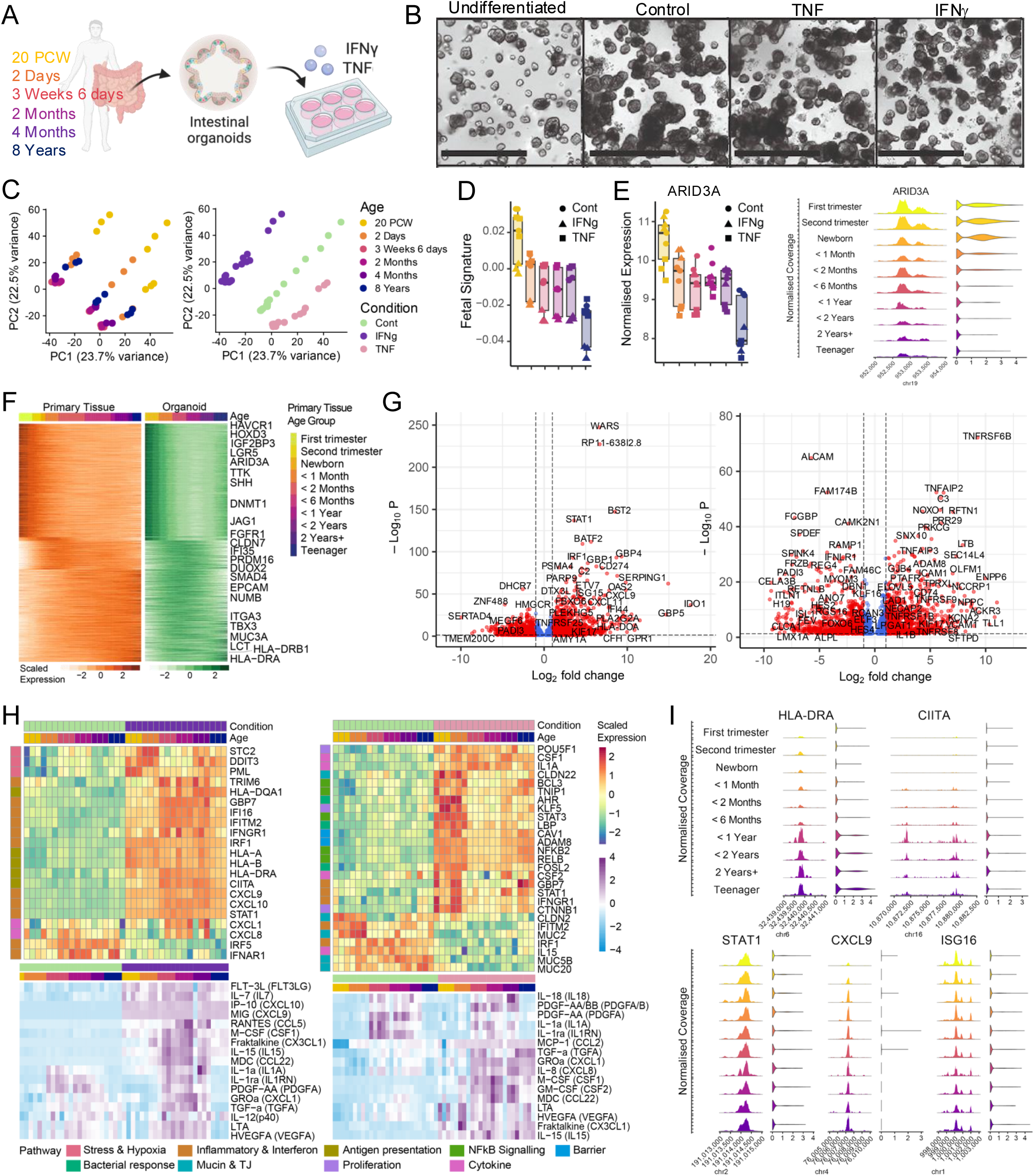
Organoids model age-dependent epithelial immune competency. A. Schematic of organoid stimulation experiments using terminal ileal organoids derived from donors of different ages and stimulated with IFNγ or TNF. Created with BioRender. B. Representative images of organoids under undifferentiated, unstimulated, TNF- and IFNγ-stimulated conditions. Scale bars, 250 μm. C. PCA of bulk RNA-seq data from organoid stimulation experiments, coloured by age and stimulation condition. D. Fetal module score (derived from ileal epithelial single cell multiomics data) projected onto organoid bulk RNA-seq samples. Points are coloured by age and shaped by stimulation condition; bar plots show distribution quantiles (Q1, median, Q3) per age group. E. ARID3A expression and chromatin accessibility. Left, scatterplot showing normalized ARID3A expression in organoids, coloured by age and shaped by stimulation condition; bar plots show distribution quantiles (Q1, median, Q3) per age group. Right, coverage and violin plots showing promoter accessibility and gene expression in primary ileal epithelium, coloured by age group. F. Comparison of gene expression dynamics between primary tissue and organoids. Heatmaps show genes significantly differentially expressed in both datasets (adjusted P < 0.05). Expression was pseudobulked (tissue), spline-smoothed, z-score normalized per gene and ordered by stage of maximal expression; columns are annotated by age group. G. Volcano plots of differential expression in IFNγ-(left) and TNF-(right) stimulated organoids relative to unstimulated controls, assessed using DESeq2 (Wald test with Benjamini–Hochberg correction). Points are coloured by significance and effect size (red, |log2FC| > 1 and −log10P > 1; blue, |log2FC| < 1 and −log10P > 1). H. Top, heatmaps showing scaled expression of selected genes with age- and stimulation-dependent changes (IFNγ, left; TNF, right). Bottom, heatmaps of Luminex protein measurements under corresponding conditions; colour scheme consistent with C. I. Coverage and violin plots showing promoter accessibility and gene expression of selected age-associated genes in primary ileal epithelial cells.

We first verified that PDOs retained age-specific transcriptional features observed in primary tissue single-cell data. Unstimulated organoids segregated by developmental age, consistent with primary tissue transcriptomes (**Figure 5C**). A fetal signature score derived from single-cell data (**Methods**) was highest in 20 PCW organoids and declined postnatally (**Figure 5D-E, Supplementary Figure 11A-C**), with fetal organoids showing increased expression of developmental genes, including *H19*, *LCN15*, *TTR*, *ARID3A* and *HAVCR1*. Differential analyses across developmental stages showed strong concordance between primary tissue pseudobulk and organoid bulk RNA-seq data (**Figure 5F, Supplementary Figure 11D-G**), with fetal and neonatal PDOs enriched for cell cycle, DNA replication and ribosome biogenesis pathways, whereas paediatric organoids were enriched for pathways related to digestion and transport, consistent with prior single-cell analyses of primary ileal epithelial cells (**Supplementary Figure 11B**).

IFNγ stimulation elicited a robust but age-dependent transcriptional response in PDOs (**Figure 5C, 5G, Supplementary Figure 12A, Supplementary Data**). Across all ages, IFNγ induced a canonical interferon program, including *STAT1*, ISGs and chemokines, together with pathways related to autophagy and immune defence (**Figure 5G**, **Supplementary Figure 12A**). However, in fetal and early neonatal organoids, some of these responses were attenuated, and instead we observed an increase in activated stress-adaptive programmes, induction of hypoxia- and cellular stress–associated genes (**Figure 5H, Supplementary Figure 12A, Supplementary Data).** Similarly, class II antigen processing and presentation pathways were strongly upregulated in response to IFNγ in organoids at three weeks of age or older, but this was reduced in fetal or early neonatal organoids (**Figure 5H, Supplementary Figure 12A**). Previous work has demonstrated that intestinal epithelial cells can present antigen to CD4⁺ T cells, with IFNγ being necessary and sufficient to induce this process in adult organoid and *in vivo* models^55^. Our data here indicates that the capacity of epithelial cells to act as antigen-presenting cells may be age-restricted. Accordingly, in primary intestinal epithelium multiome data, the HLA-DRA promoter was not accessible in fetal or early paediatric samples but a strong peak was detected only 6 months postnatally (**Figure 5I).** IFNγ-induced MHC-II expression is mediated by the master regulator CIITA, whose IFNγ-responsive promoter contains STAT1 and IRF1 binding elements^56^. We observed a strong postnatal gain in chromatin accessibility at the promoter upstream of the CIITA exon, as well as at intronic regulatory elements within the CIITA locus (**Figure 5I**). Examining all IFNγ-induced genes in older pediatric but not fetal or early post-natal organoids, we observed that 53% (632 of out 1195) of DEGs showed age-increasing promoter and/or enhancer accessibility in primary epithelium (**Supplementary Figure 12B-C**). Together, this suggests that the chromatin landscape of early life epithelium is not permissive for full induction of IFNγ responses and that increasing epigenetic “licencing” during maturation may be a key mechanism for modulating inflammatory signalling. Recent work in fetal epithelial organoids^57^ has demonstrated that IFNγ from fetal T-cells can promote barrier maturation, further supporting developmental-stage specific roles for IFNγ-epithelial signalling.

Whereas IFNγ stimulation activates JAK-STAT1 and induction of MHC I and II, TNF stimulation induces NF-κB–driven cytokine programmes that affect epithelial survival and barrier function. Overall, while TNF elicited less pronounced changes relative to control than IFNγ (**Figure 5G-H, Supplementary Figure 12A, Supplementary Data**), stimulation robustly activated NF-κB signalling (*TNFAIP2*, *NFKB1*, *IL1B*) across all age groups. On the other hand, age-dependent effects included a group of fetal and early neonatal-expressed genes (**Supplementary Figure 12A,** Cluster C1) encompassing development and morphogenesis functions that were suppressed by TNF stimulation, suggesting that cytokine stimulation may accelerate aspects of epithelial maturation. A further group of genes encompassing response to metal ion, organic acid transport and response to toxic substance (**Supplementary Figure 12A,** Cluster C2), were induced in older but not fetal or early pediatric organoids. We further identified a gene module that was more strongly induced in fetal and neonatal organoids than at later stages in response to TNF stimulation (**Supplementary Figure 12A,** Cluster C3). This module was enriched for regulators of NF-κB and cytokine signalling (*TNFRSF1A, NFKB2, RELB, BCL3, TNIP1, STAT3*), epithelial proliferation and Wnt/β-catenin activity (*KLF5, CTNNB1, POU5F1*) and barrier- and membrane-associated pathways (*AHR, CAV1, CAV2, ADAM8*), suggesting a heightened ability of early life epithelium to promote barrier integrity and epithelial proliferation in response to TNF. Similarly to age-dependant IFNγ responses, we found that a substantial fraction of pediatric-only DEGs could be explained by increasing promoter or enhancer accessibility in epithelium (**Supplementary Figure 12B-C**). In line with our findings across fetal and postnatal organoids, it has been previously demonstrated that TNF can promote growth in fetal epithelial organoids in a dose-dependent manner ^58,63^.

In order to assess whether these transcriptional changes also corresponded to cytokine secretion, we undertook multiplex bead-based immunoassay (Luminex) quantification of epithelial secreted cytokines from PDO supernatants. We observed increased TNF-induced secretion of CXCL1, CXCL8, CSF1, CSF2 and PDGFA/B in post-neonatal and older organoids, but not fetal and neonatal organoids (**Figure 5H**). Similarly, IFNγ stimulation led to increased secretion of CXCL1, CXCL8, IL1A, CCL22 and IL15 (**Figure 5H**) in post-neonatal, but not fetal or neonatal organoids. Conversely, CXCL9, CXCL10 and IL7 cytokine secretion was uniformly induced across all age groups. Consistent with these findings, single cell data revealed chromatin at CXCL9 locus, for instance, was accessible from second trimester (**Figure 5I, Supplementary Figure 6B**), while CXCL8 showed transient accessibility later (**Supplementary Figure 12C**). Together, these data indicate that early-life epithelium may not be epigenetically “licenced” to elicit a full range of pro-inflammatory cytokine-induced innate immune responses, but this window is very brief.

### Postnatal BHLHE40 attenuates epithelial inflammatory responses

We next sought to use organoids to explore how these post-natal epithelial inflammatory responses are attenuated during the transition to infancy. Our previous analyses revealed early-life TF regulatory networks that likely enable epithelial adaptation to rapid environmental changes (**Figure 2F**). Among these, BHLHE40 was implicated as a potential regulator of inflammatory programs in the early postnatal period (**Figure 3A-B, 6A, Supplementary Figure 8B**), with predicted targets enriched for antiviral and antibacterial responses, interferon signalling and antigen presentation (**Figure 6B**). BHLHE40 functions predominantly as a transcriptional repressor by binding E-box motifs and recruiting HDAC-containing corepressor complexes, although in specific contexts, such as stress responses or metabolic regulation, it has been reported to indirectly promote transcription ^59–61^. In our single cell multiome cohort, we observed BHLHE40 expressed in fetal epithelium, followed by a peak in early neonatal samples and a subsequent decrease in older infants (**Figure 6A**). IHC and ISH analysis of an independent tissue cohort in ileum confirmed robust epithelial expression of BHLHE40, with higher levels in fetal and early postnatal tissue (**Figure 6C, Supplementary Figure 13A**).

**Figure 6.**
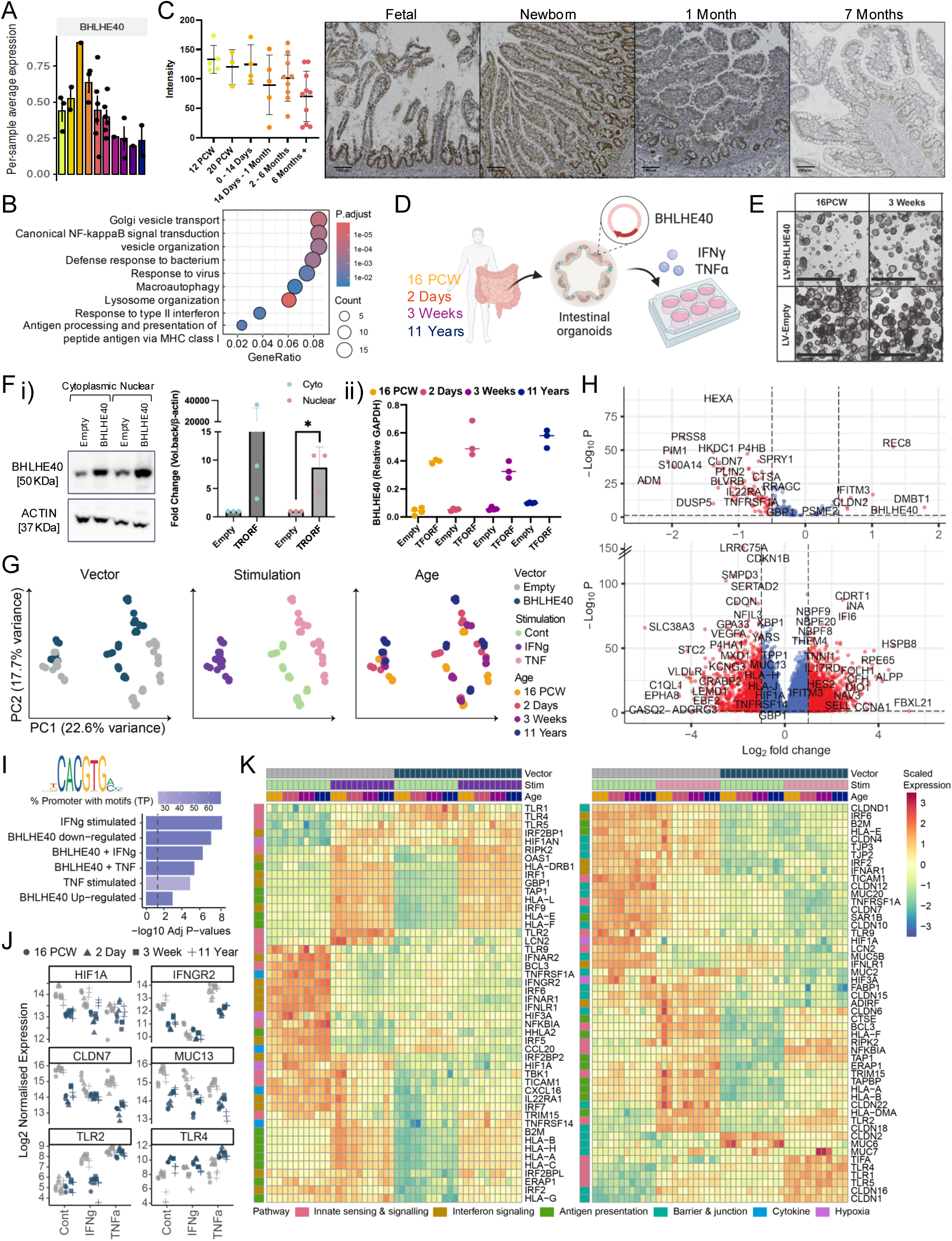
BHLHE40 attenuates epithelial inflammatory responses. A. Mean expression of BHLHE40 in ileal epithelial cells from single cell multiomics primary tissue data across age groups; points represent individual donors. B. Dot plot of pathways enriched among BHLHE40 target genes. C. Protein-level validation of BHLHE40 expression across development. Left, quantification of in situ hybridization (ISH) staining; points represent donors. Right, representative immunohistochemistry (IHC) images across developmental stages. Scale bars, 100 μm. D. Schematic of organoid experiments with lentiviral overexpression of BHLHE40 followed by IFNγ or TNF stimulation. Created with BioRender. E. Representative images of organoids transduced with control or BHLHE40-expressing vectors. Scale bar, 250 μm. F. Validation of BHLHE40 overexpression. i) Western blot of 2-day organoids and quantification of protein levels. ii) qPCR quantification of BHLHE40 expression. G. PCA of bulk RNA-seq data from organoids with BHLHE40 overexpression and cytokine stimulation, coloured by age, vector and stimulation condition. H. Volcano plots showing differential expression in BHLHE40-overexpressing organoids compared to controls (top, known targets; bottom, all genes), assessed using DESeq2 with Benjamini–Hochberg correction. Points are coloured by significance and effect size. I. Enrichment of BHLHE40 binding motifs in promoters of differentially expressed genes following BHLHE40 overexpression and/or cytokine stimulation, assessed relative to non-differentially expressed genes. J. Scatterplots showing normalized expression of selected genes in organoid RNA-seq data, coloured by vector (same colour scheme as G). K. Heatmaps showing scaled expression of selected genes across conditions, with annotations for vector, stimulation, age and pathway classification; colour scheme consistent with G.

In order to assess the function of BHLHE40 in early-life inflammatory responses in epithelium, we overexpressed BHLHE40 in intestinal organoids via lentiviral transduction and subsequently stimulated them with TNF and IFNγ(**Figure 6D-E, Supplementary Figure 13B**). Successful Lv-BHLHE40 over-expression was confirmed at mRNA and protein level by qPCR, sequencing and Western blot respectively (**Figure 6F, Supplementary Figure 13C-D**) and downstream transcriptional changes were assessed by bulk RNA-seq. Principal component analysis confirmed that BHLHE40 overexpression accounted for the second largest sources of variance within this data, after the effects of cytokine stimulation, indicating robust downstream effects across all organoid lines (**Figure 6G**).

Differential expression analysis identified 6299 upregulated and 5804 downregulated genes in BHLHE40 overexpressing organoids (**Figure 6H**). At baseline in unstimulated organoids, BHLHE40 increased expression of DNA replication, RNA processing and retinoic acid metabolism genes, including stem cell marker *ASCL2*, and attenuated transport, hypoxia response and both class I and class II antigen presentation pathways. Motif analysis of DEG promoter regions identified a significant enrichment of BHLHE40 binding motif in the promoter regions of significantly downregulated, but less in upregulated genes, consistent with its canonical repressive role (**Figure 6I**); therefore, it is likely that upregulated gene expression may arise indirectly as a result of, for example repression of other transcriptional repressors.

Notably, BHLHE40 overexpression increased expression of *TLR4*, suggesting a redirection of epithelial innate sensing toward TLR4-mediated pathways to facilitate microbial sensing and trigger antimicrobial peptide production^62^(**Figure 6J-K**). In addition, BHLHE40 induced expression of *CLDN2*, a marker associated with increased epithelial permeability, while TNF stimulation further enhanced expression of *CLDN1*, *CLDN16*, and *TLR6* (**Figure 6J-K**). These context-dependent effects suggest that BHLHE40 does not globally suppress epithelial defence but instead reshapes barrier architecture and innate sensing capacity in response to inflammatory cues.

Upon stimulation with IFNγ and TNF, we observed robust induction of canonical interferon and NF-κB signalling programs respectively in control organoids, replicating our previous experiments (**Supplementary Figure 13E, Supplementary Data**, 13045 and 9285 DEGs respectively); however, BHLHE40 overexpressing organoids exhibited a reduced response. Differential gene expression analysis identified 4365 genes that were upregulated by IFNγ in control organoids with reduced expression in BHLHE40 overexpressing organoids and 2072 genes following this pattern upon TNF stimulation (**Figure 6K, Supplementary Data, Supplementary Figure 13F).** In IFNγ, these encompassed interferon-stimulated genes involved in antiviral defence (including IRF1, IFITM1/2/3, OAS family members and GBP genes), components of antigen processing and presentation machinery (HLA class I and II molecules, B2M, TAP1 and ERAP1), mediators of innate inflammatory signalling (NFKBIA, TNFRSF1A, RIPK2 and TBK1) and stress-response programmes. TNF non-response genes comprised core components of canonical NF-κB signalling (e.g. TBK1, NFKBIA, TNFRSF1A, BCL3, TIFA), antigen processing and MHC class I presentation machinery (TAP1, TAPBP, ERAP1, B2M, HLA-A/B/E/F, CTSE, SAR1B), barrier and epithelial differentiation programmes (MUC2, FABP1, CLDN22) and innate immune sensing pathways, together indicating that BHLHE40 overexpression attenuates epithelial inflammatory programs downstream of TNF and IFNγ stimulation.

## Discussion

We provide a comprehensive single-cell and spatial atlas across fetal, neonatal and paediatric development, constructed with single cell multi-omics and MERSCOPE spatial transcriptomics. Using these data, we generated a multimodal molecular clock and a deep learning-based transcriptomic model to chart early intestinal maturation. Our analysis highlights transient postnatal cell populations and molecular programmes that drive epithelial barrier maturation and immune homeostasis. We define the regulatory networks underpinning post-natal epithelial imprinting, junctional expression dynamics, and spatial niches shaping neonatal epithelial defence. Functional modelling with intestinal organoids revealed BHLHE40 as a key early-life regulator of epithelial inflammatory programmes. Collectively, these findings clarify the pattern of epithelial stem cell imprinting at specific developmental milestones; the principles of epithelial remodelling governing post-natal sealing and innate reactivity and the cellular dynamics linked to niche establishment in early life. This atlas provides a foundational reference for human intestinal development and a framework for understanding developmental disorders and early-life diseases such as necrotising enterocolitis.

Our study highlights the immediate postnatal period as a brief, molecularly distinct phase of coordinated epithelial, stromal and immune remodelling. Rather than a uniform “immaturity,” early-life epithelium exhibits stage-specific functional programmes: differentiated fetal enterocytes retain distinct transcriptional identities that rapidly reprogramme after birth to support lipid metabolism, xenobiotic handling and localized innate defence. Chromatin profiles show that regulatory readiness (open promoters/enhancers) often precedes transcriptional activation, indicating epigenetic priming that enables rapid postnatal adaptation. Transient retention of fetal regulons (e.g., ARID3A-associated networks) in neonates resolves within the first weeks, implying a short - but not immediate-transitional window after birth. Conversely, a subset of fetal stem cell-associated regulatory elements remains accessible after birth despite reduced transcriptional activity. This persistent accessibility across both stem and differentiated epithelial lineages suggests a retained epigenetic competence that may enable rapid reactivation of regenerative and barrier-adaptive programmes when required.

Development proceeds asynchronously across compartments. Epithelial, myeloid and T-cell programmes change earliest, while B/plasma cells and multiple fibroblast states continue to evolve through infancy. This asynchrony likely reflects differing physiological functions: the epithelium must quickly adapt to achieve nutrient uptake and symbiose with the microbiota, whereas adaptive immunity and tertiary lymphoid structures continue to adapt to establish intestinal adaptive immune protection. Fibroblasts must continue supporting intestinal growth and expansion of their specialised niches later into childhood.

We refine the “window of opportunity” concept by showing it is region- and gene-specific: tight-junction and polarity components follow heterogeneous trajectories (e.g., CLDN2 dynamics differ between ileum and colon) and sealing claudins are not uniformly downregulated at birth. Junctional dynamics are complex around birth and it will be interesting to assess the functional relevance of these changes in future work. For example, it will be interesting to explore the temporal role of selective downregulation of sealing claudins in paracellular flow, microbial handling, M cell biology and antigen presentation and the promotion of an immune regulatory landscape post-birth.

Epithelial innate defence programmes are not maximally induced immediately after birth. DUOX2 and associated ROS and innate defence pathways increased after the first postnatal week, peak within the first month and subsequently decline. This timing aligned with our organoid stimulation experiments. The earliest fetal and newborn organoids exhibit attenuated cytokine responsiveness, while organoids derived after the first weeks of life demonstrate more robust interferon and antigen-presentation responses. Together, these findings support a model in which the earliest epithelium is epigenetically constrained and only partially licensed for inflammatory activation. This staged responsiveness likely protects the stem-cell niche while permitting progressive immune education.

Mechanistically, we identify BHLHE40 as an epithelial transcriptional regulator that likely modulates early life epithelial immune responses. BHLHE40 overexpression in organoids dampens antigen-presentation, hypoxia and NF-κB–related programmes and represses many predicted targets, indicating a role in limiting epithelial pro-inflammatory signalling. Interestingly, IFNγ but not TNF stimulation downregulated endogenous BHLHE40 expression (**Supplementary Figure 13G**), suggesting that BHLHE40-mediated suppression of epithelial responses can be overridden by specific stimuli. In our data, this coincides with the shifting cytokine milieu within the first few weeks of life - Th1 cell numbers are low at birth, but myeloid cells capable of producing TNF are more prevalent, and BHLHE40 expression by epithelium declines with emergence of a more diversified T-cell and IEL compartment. Consistent with this, IHC demonstrated that BHLHE40 expression becomes progressively restricted to the crypt base in older epithelium (**Figure 6C**), suggesting that its inflammation-attenuating effects may act within the stem cell niche to protect genomic integrity during ongoing epithelial renewal. The exact stimuli leading to induction of BHLHE40 in the post-natal period remain to be determined but in other contexts it can be induced by retinoic acid (RA) or IL1^60^. Colonisation with RA expressing microbes may drive enhanced epithelial BHLHE40 activity and barrier immunoregulation.

It is worth noting that while our analyses characterised transient pro-inflammatory immune cell signalling in early life, these activated cell states were confined to epithelial niches constituting a minority of overall mucosal immune cell populations. Most macrophages retained M2-like transcriptional profiles, with only a minor subset adopting M1-associated signatures. Similarly, although cytotoxic and activated T-cell programmes peaked within the first postnatal month, the majority of T cells were naïve, memory or regulatory in phenotype rather than effector-dominant. Furthermore, all samples analysed were histologically normal and non-inflamed. Thus, the early-life cytokine milieu likely reflects a developmentally programmed, low-grade inflammatory alert state that arises due to first encounters with microbes and supports immune and barrier education rather than overt inflammation.

### Limitations of the study

Currently, our patient-derived organoids successfully recapitulate core fetal, neonatal, and older postnatal epithelial features. However, some transient and age-restricted gene expression programmes identified in primary cells are attenuated or lost in culture. Beyond the isolated epithelium, there are open challenges in faithfully modelling fine-grained developmental molecular milestones without the inclusion of stromal, immune, and microbial inputs.

Here, we have established a predominantly repressive role for BHLHE40. However, given that BHLHE40 function can be context-dependent (with some predicted target genes increasing upon overexpression), the identification of precise regulatory mechanisms may not be a case of simple extrapolation. We currently lack direct binding data (e.g., ChIP-seq) and therefore cannot definitively distinguish direct from secondary, network-level transcriptional effects.

Further important future directions therefore include (i) expanding age-stratified organoid lines and developing complex co-culture systems to reconstruct epithelial maturation, and (ii) utilizing chromatin-based analyses to clarify the mechanisms underlying BHLHE40-mediated regulation.

We have successfully identified core age-specific epithelial expression programmes and key regulatory mechanisms during postnatal development. These results lay the foundation for more complex, multi-cellular models of infant intestinal maturation, promising advancements in our understanding of paediatric intestinal health and disease.

## Acknowledgements

Funding for this study was generously provided by grant number DAF2021-237606 from the Chan Zuckerberg Initiative DAF, an advised fund of Silicon Valley Community Foundation (or the Chan Zuckerberg Initiative Foundation); a Wellcome Career Development Award (315684/Z/24/Z) to A. An.; NIHR Academic Clinical Lectureship to D.F.-C; The Academy of Medical Sciences PPI grant to D.F-C; University of Oxford Medical Sciences Internal Fund (0018327) to D.F-C; Swiss National Science Foundation (P500PB_225639) grant to V.L; an NIHR Senior Investigator Award (NIHR201410); a Wellcome Investigator Award (219523/Z/19/Z) (A.S.); the UK Medical Research Council MC_UU_00036/1 (A.S.). C.M. was also supported by Oxfordshire Health Services Research Committee (OHSRC, ref. 1407), part of Oxford Hospitals Charity.

This study was made possible through the collaborative efforts of the MIMIC Consortium clinical teams across multiple sites. Oxford Children’s Hospital: The paediatric surgery team including Khaled Ashour, Eva Coates, Alex Lee, Kokila Lakhoo, Merill McHoney, Ibrahim Mostafa, Ashok Rijhwani, and Ian Willetts. Research support was provided by Frey Carvalho, Sarah Clews, Leah Varghese and the Children’s Research Team; Southampton Children’s Hospital: The paediatric surgery team including Charles Keys, Lara Kitteringham, Ori Ron, Michael Stanton, Francesca Stedman, and Robert Wheeler. Research support was provided by Gavin Babbage, Jevan Hales, Irina Ferreria Fino, Jenny Pond, Amy Whiteman, the Children’s Research Team and the Clinical Research Facility. Alder Hey Children’s Hospital: The paediatric surgery team including Sarah Almond, Simon Kenny and Josef Taylor. Research support was provided by the Children’s Research Team.

Further patient recruitment and study co-ordination was undertaken by ORNIID research nurses and Oxford TGLU Biobank, led by S. Fourie and J. Chivenga, respectively - we thank them, the patients and their parents who made this work possible.

Human fetal material was provided by the Joint MRC/Wellcome Trust (MR/R006237/1) Human Developmental Biology Resource (https://www.hdbr.org), and we thank B. Crespo, Professor Copp, and the entire HDBR team.

We thank Dr. Karina Pombo-Garcia and Dimitrios Ioannidis for ongoing discussions on epithelial tight junctions.

We thank parents and families from the Parent Neonatal Advisory Group for their guidance, perspective and dedication to improving care for children. We are deeply grateful to the patients and parents who participated in the MIMIC study. We thank them for their generosity in contributing samples, without which this research would not have been possible.

## Author Contributions

A.S, A.An, D.F-C and C-H-J.L designed the study. C.H-J.L, A.An, D. F-C, A.S and V.L wrote the manuscript. C.H-J.L, A. An, P. Z-S. and V.W.L undertook multiome data analysis. C.H-J.L and A. An undertook spatial transcriptomics data analysis. C.H-J.L and A. An undertook bulk RNA-Seq data analysis. C.H-J.L and D.F-C undertook Luminex data analysis. C.H-J.L developed molecular clock model. A.An developed data portal. D.F.C, L.S, V.L, A.G and M.J undertook organoid experiments. L.S and L.D undertook western blot experiments. L.S, V.L, D.F-C and A.M undertook IHC experiments. L.S and D.F-C undertook ISH experiments. D.F-C and L.S undertook qPCR experiments. D.F-C and L.S undertook Luminex experiments. Z.C, A.S-G, D.F-C and P.Z undertook single cell multiomics experiments. Z.C, E.B, R.K, P. V.G, C.C, M.G and C.M undertook spatial transcriptomics experiments. D.F-C, L.S, X.Q, H-W.C and M.J undertook lentiviral transduction experiments. Z.C, A.S-G, V.L, V.W-L, L.S, P. G-C, A. Au, N.C, T.G and C.M undertook the processing of biological samples and contributed significant technical expertise to the development and refinement of experiments. S.B. provided significant input into experimental design and execution. D.F. provided significant histopathology input. G.R, R.H, N.J.H and P.J led MIMIC patient recruitment and provided clinical input. H.K provided additional senior oversight. A.S. and A.An co-supervised.

## Methods

## EXPERIMENTAL MODELS AND SUBJECT DETAILS

### Sample Collection, Handling and Processing

Samples were taken during clinically mandated visits, in compliance with local and national patient safety directives, and with informed consent (MIMIC Study, Ethics-22/NS/0027; TIP-18/WM/0237; GI Biobank-16/YH/0247). Pediatric (0-18 years) samples were obtained in children undergoing intestinal resection as part of their clinical care for a range of intestinal pathologies, with sampling from the relatively healthy resection margins in tissue which was in excess to histopathological requirements. Tissue samples were transported on ice in DMEM supplemented with 10% fetal calf serum (FCS, Sigma-Aldrich), 100 U/mL penicillin, and 100 µg/mL streptomycin (Sigma-Aldrich) and processed immediately with division into formalin fixation, or sectioning into small pieces and cryopreservation as previously described ^1^.

Tissues from fetal intestine were initially collected and processed at the Human Developmental Biology Resource (HDBR), London. Following written informed consent and anonymization (HBDR project 200659, REC: 18/LO/0822), samples were collected, and intestinal tissue was dissected or isolated. The intestinal tissue was placed in Leibovitz medium (L-15, Sigma) on ice and immediately transferred to Oxford. Transported pediatric and fetal samples were washed in phosphate-buffered saline (PBS) and either frozen immediately in CryoStor® cell cryopreservation media (CS10, Sigma-Aldrich) for downstream experiments or fixed in formalin as previously described^1^.

For Formalin-Fixed Paraffin-Embedded (FFPE) sections, tissue samples were fixed in formalin for 24-48 hours before being transferred to 70% ethanol and embedded in wax through a standard graded series.

## METHOD DETAILS

### Single-cell multiome sequencing Cell isolation and staining

Tissue dissociation was performed as previously described^2–4^. Briefly, for enrichment of the epithelial fraction, intestinal resections were thawed in a 37 °C water bath. The tissue pieces were then agitated at 37 °C in Hank’s balanced salt solution (HBSS, Gibco), supplemented with 1 U/mL penicillin and streptomycin, 1% FCS, 5 mM ethylenediaminetetraacetic acid (EDTA, Invitrogen), and 2 mM dithiothreitol (DTT, ThermoFisher). The supernatant, containing epithelial crypts, was digested into a single-cell suspension using TrypLE™ Express Enzyme (ThermoFisher) for 35 minutes at 37 °C. After digestion, cells were washed, and dead cells and debris were removed using a 70-micron filter. During epithelial crypt dissociation the remaining tissue fraction, depleted for epithelial crypts and containing lamina propria cells, was dissociated using the Lamina Propria Digestion Kit (Miltenyi) for 45 minutes with regular mechanical agitation through mixing or a blunt needle. After digestion, cells (epithelial enriched and depleted fractions) were washed in Phosphate buffered saline (PBS, Sigma-Aldrich) supplemented with 1% FCS and 5mM EDTA, and dead cells and debris were removed using a 70-micron filter. Cells were then resuspended in PBS supplemented with 2% bovine serum albumin (BSA, Sigma-Aldrich) and 0.01% Tween-20 (Bio-Rad). Cells’ Fc receptors were blocked for 10 minutes with TruStain FcX (BioLegend), and cells from each participant were uniquely barcoded with TotalSeq-A hashing antibodies (BioLegend). Cells from multiomics experiments were further stained with TotalSeq™-A Human Universal Cocktail, V1.0 (BioLegend). Lamina propria-derived cells were further stained with flow cytometry antibodies for 30 minutes at 4°C. To sort stromal and immune cells anti-CD45, anti-EpCAM, anti-CD16, and anti-CD15 were used, with DAPI (4′,6-diamidino-2-phenylindole, BD) for live/dead differentiation prior to sorting. After staining cells were washed three times with staining buffer (PBS with 2% BSA and 0.01%Tween-20) before proceeding to a flow cytometer sorter (BD FACS Aria IIIu, BD FACS Fusion). EpCAM-CD16-CD15-cells were sorted into CD45+ and CD45-populations to enrich for immune and stromal cells, respectively (**Supplementary Figure 1A**).

### Epithelial cells, CD45^+^ and CD45^-^ cells

Epithelial cells and lamina propria cells were isolated and stained as described above. Cells from intestinal samples of 4-5 patients were pooled together from each compartment (epithelial, CD45+, and CD45-). Compartmental cells (epithelial, CD45+, and CD45-) were all processed following the DOGMA-seq protocol^5^ and the 10X Genomics protocol (10X Genomics, CG000338, Rev F) with modifications. Briefly, cells were spun down, and the pellet was resuspended in 100 µL of cold digitonin lysis buffer (0.01% Digitonin, 20 mM Tris-HCl pH 7.5, 150 mM NaCl, 3 mM MgCl₂, and 2 U/µL RNase inhibitor). Pooled epithelial cells were incubated on ice for 7 minutes, and CD45+ and CD45-compartments for 3 minutes. At the end of the incubation, cells were washed in 1 mL of digitonin wash buffer (20 mM Tris-HCl pH 7.5, 150 mM NaCl, 3 mM MgCl₂, and 1 U/µL RNase inhibitor) at 500xg for 5 minutes at 4°C. Cells were then resuspended in 150 µL of digitonin wash buffer and counted using Trypan blue staining to verify permeabilization (Countess 2, Thermo Fisher). For each run, an input of 30,000-40,000 single cells per pool was added to each channel of the Chromium X single-cell platform (10X Genomics) with a recovery rate of approximately 10,000 cells.

### Library construction and sequencing

To further enhance sequencing quality, the Jumpcode Deplete X kit was used according to the manufacturer’s instructions. The kit leverages Cas9 depletion with an optimized guide set to remove ribosomal and mitochondrial mRNA, as well as non-transcriptomic reads. The rest of the library construction was carried out following the Chromium Next GEM Single Cell Multiome ATAC + Gene Expression protocol (10X Genomics, GC000338 Rev F), incorporating the suggested modifications of the DOGMA-seq protocol.

Specific protocol modifications and all primers used for surface library construction can be found in custom oligos table (**Supplementary Data**). The quality of RNA, ATAC, and HTO libraries was assessed using the Bioanalyzer High Sensitivity DNA kit (Agilent Technologies), and concentrations were measured with the Qubit dsDNA HS Assay Kit (Invitrogen). Sequencing of the libraries was performed on either an Illumina NextSeq 500, a NovaSeq 6000, or a NovaSeq X Plus platform, with a target of ∼80,000 ATAC reads per cell, ∼80,000 RNA reads per cell, and ∼10,000 HTO barcode reads per cell.

### Spatial Transcriptomics - MERSCOPE

RNA was extracted from samples using the Deparaffinization Solution (Qiagen) and the RNeasy FFPE Kit (Qiagen). RNA integrity and quality were assessed using a High Sensitivity RNA ScreenTape assay and a 4200 TapeStation (Agilent Technologies). The DV200 of all samples was greater than 50%.

The MERSCOPE Vizgen Platform was used for high-resolution subcellular spatial transcriptomics, following the manufacturer’s instructions for resistant tissue with modifications (Vizgen, MERSCOPE, 91600112, Rev B). Briefly, paediatric and fetal FFPE tissues were microtome sectioned at a thickness of 5 µm and placed on MERSCOPE slides. Slides were incubated in an oven at 55°C for 15 minutes and then air-dried for 2 hours before being stored at −20°C for a maximum of 3 days. Subsequently, sections underwent deparaffinization, cell boundary staining, anchoring, gel embedding, digestion for 6 hours, and extended clearing. Quenching was performed for 6 hours to limit autofluorescence before probe hybridization. The slides were incubated with custom probes (**Supplementary Table**) for 68 hours prior to imaging on the MERSCOPE platform, according to the manufacturer’s instructions (Vizgen, 91600001, Rev F).

### Organoid stimulation with pro-inflammatory cytokines

To establish organoid lines, tissue was collected form the terminal ileum of fetal and paediatric patients (macroscopically non-inflamed). Epithelial crypts were isolated using EDTA chelation as previously described. In brief, previously dissected (<0.2cm^2^) cryopreserved tissue was thawed for 2 minutes at 37° in a water bath. Tissue was washed in 10ml of HBSS supplemented with 1% FCS. Epithelial crypts were chelated from tissue in 3 x 10 minute incubations at 37°C in 10ml chelation media (HBSS supplemented with 100u/ml penicillin/0.1mg/ml streptomycin; 10mM HEPES, 5mM EDTA and 1% Fetal Bovine Serum) with agitation every 3-4 minutes to isolate crypt fraction in supernatant. Supernatants from the 3 rounds were pooled and and resuspended in WREN medium (WNT3A conditioned media 50%, Advanced DMEM-F12 media, 2mM glutamax, 10mM HEPES, N2 supplement, B27 supplement, 10mM nicotinamide, 1mM N-acetyl-L-cystine, 50ng/ml human EGF, 1µg/ml gastrin, 0.1µM A83-01, 10 µM p38 inhibitor SB202190, 10% Noggin conditioned media and 10% R-Spondin conditioned media then plated in Matrigel (Corning) at a 1:1 ratio. 10 µM Y27632 was added at isolation and passaging ^6^. Media was changed every 2-3 days until organoids were generated and passaged every 7-10 days by breaking of domes with cold PBS, dissociation with TrypLE express (3 minutes, 37 °C), filtering and re-culturing in Matrigel and WREN media.

For stimulation experiments organoids were generated from a timeline of ages (**Supplementary Data**) and grown to passage 4, then established in 24 well plates and grown in 400ul WREN media for 6 days, and continued in WREN supplemented with respective cytokines (control, IFNγ or TNF-α at 100ng/ul) for an additional 2 days before differentiation for 3 days in Intesticult Organoid differentiation media (StemCell) with respective cytokine. On day 11 differentiated organoid domes were broken down with cold PBS (4°C) and RNA isolated using QIAshredder and RNeasy Mini kit (Qiagen) as per manufacturer’s instructions. Two Matrigel domes per condition were pooled for RNA extraction and experiments were performed in quadruplicate. Supernatants were harvested and stored at −80°C until downstream analysis.

Eluted RNA was quantified with spectrophotometry (Nanodrop) and quality control performed with Agilent 4200 Tapestation using High Sensitivity RNA Screentape to confirm RNA integrity (eRIN >9.0). Following QC RNA libraries were indexed and pooled for sequencing on Novoseq X plus (150PE reads, Illumina) aiming for 6Gb reads per sample.

Supernatants were thawed and processed on a multiplex bead immunoassay as per manufacturer’s instructions, with custom panel bead array (Merk Millipore, HCYTA-60 Human Cyto panel A, 27 targets, **Supplementary Data**). Following staining and processing beads were quantified on INTELLIFLEX instrument (Luminex).

### Lentiviral production and transduction of intestinal organoids

Intestinal epithelial organoid lines were established as previously described. Lentiviral expression vector for *BHLHE40* were obtained from F. Zhang^7^ via Addgene (TFORF2593). The empty vector was generated by excising the insert from the lentiviral vector using sing NheI-HF and SpeI-HF restriction enzymes (New England Biolabs), followed by gel purification and self-ligation of the linearized vector. HEK293 cells were obtained from ATCC (CRL-1573) and tested to be mycoplasma-free during culture and passage, maintained with DMEM supplemented with 10% FBS and 1x penicillin/streptomycin in culture. HEK293 cells were seeded and at 90% confluence in T25 flasks, then transfected with 2.6µg of psPAX2 (Addgene), and 1.7 µg of pMD2.G (Addgene 12259); alongside 3.4 µg of TFORF2593 or empty vector respectively, in the presence of FuGENE HD transfection reagent (Promega) at a ratio of 3:1 to vector DNA. Viral supernatant was collected at 48 hour, and 96 hours with supernatants filtered through a 0.45-µm PVDF membrane, aliquoted, and stored at –80 °C.

For transduction, intestinal organoid cultures (passage 3-4) from primary tissue across ages (16pcw, 2 days, 3 weeks, 11 years) were harvested, dissociated in TrypLE express (5-10 minutes, 37 °C) then spinoculated in a 24 well plate (100,000 live cells per well, 600xG 1 hour room temperature) with organoid basal transfection media (DMEM/F12 supplemented with HEPES, Glutamax, B27 and N2) and lentiviral supernatant at optimised MOI with addition of 0.8ug/ml polybrene (Santa Cruz). Spinoculated cells were then incubated for a further 5 hours at 37 °C, collected and cultured in WREN with antibiotics omitted. 48 hours later, after small cystic organoids were forming, puromycin (1µg/ml) was supplemented to media to undertake 6 days of puromycin selection, then organoids were passaged, and cultured as previously described. For stimulation experiments transfected organoids (TFORF2593 and empty vector) were cultured, stimulated and differentiated in a manner previously described for non-transfected organoids with RNA extracted and mRNA sequencing performed.

### Real time PCR

For RNA quantification of *BHLHE40* expression in transfected organoids, organoids were grown, transfected and once confluent total RNA extracted as previously described (**Organoid stimulation with pro-inflammatory cytokines)**. RNA was reverse transcribed to cDNA with the High-capacity cDNA reverse transcription kit (Applied biosystems) as per manufacturers instruction with maximum 1000ng input, quantified by Nanodrop, and final cDNA diluted to 5ng/µl. For quantitative RT-PCR samples were then measured on the Quantstudio 7-flex system (ThermoFisher) with applicable Taqman gene expression assays and quantified to housekeeping (*GAPDH*) with experiments performed in at least biological and technical triplicates and analysed with the ddCT method.

### Western Blot

Cytoplasmic and Nuclear fractions were obtained from 1×106 cells following the protocol developed by Huynh et al ^8^. In brief, HLB and NLB buffer supplemented with protease and phosphatase inhibitors were used to separate the cellular fractions prior to sonication for 15 seconds at 60% amplitude. 10 µg of protein was loaded to precast BoltTM 4-12%, Bis-Tris Plus WedgeWellTM Gels (Thermo Fisher Scientific). Proteins were transferred to a PVDF membrane using the iBlot 2. After blocking in non-fat milk, membranes were incubated overnight at 4 °C with primary BHLHE40 antibody (1:500). Membranes were subsequently incubated with secondary antibody at 1:1000 (Anti-Rabbit IgG) and developed using SuperSignal™ West Pico PLUS Chemiluminescent Substrate prior to imaging on the ChemiDoc MP System. After imaging the membrane was stripped with RestoreTM Stripping Buffer to blot other proteins of interest. β-Actin was used as a reference protein, at a concentration of 1:2000. Band intensities were quantified using the iBright Analysis Software and normalised to β-Actin.

### Immunohistochemistry

Paraffin-embedded tissue sections were microtome sectioned at 4μm and mounted on Superfrost slides. Slides were deparaffinised in Histoclear and rehydrated using an ethanol series. Heat induced epitope retrieval was performed at 96 °C in either pH6 or pH9 Tris/EDTA buffer for 30 minutes. Endogenous peroxidase was blocked using H2O2 before slides were blocked using 2% goat serum for 1 hour. Slides were incubated with primary antibody (listed below) diluted in 1% BSA overnight at 4 °C. Post washing, samples were developed with appropriate species specific ImmPACT HRP kits (anti-rabbit or anti-goat) according to manufacturer’s instructions prior to developing in DAB and counterstaining with Mayer’s Haematoxylin. Slides were dehydrated using an ethanol series and mounted using Leica Mounting Medium. High resolution 20X images were taken using the Zeiss Axio-Scan Z1.

Scans were imported into Visiopharm (version 2025 08.6 x64), tissue sections were identified using the in-built Tissue Detection app. Bubbles and tissue folds were identified using a custom built deep-learning app and excluded from analysis. Post tissue clean-up; crypts were identified and segmented using a custom-built threshold algorithm. Following crypt segmentation, colour deconvolution was used to measure DAB intensity, providing a quantification of the staining intensity.

### Single molecule in situ hybridisation (ISH)

Paraffin embedded samples were sectioned in the same manner as IHC, and pre-heated to 60 °C for one hour prior to use. Slides were deparaffinised in Xylene and rehydrated using an ethanol series. Probes and development reagents were obtained from Advanced Cell Diagnostics (ACD) RNAscope® kits. smISH was performed as per manufacturer’s instructions (322310-USM), using a HybEZ oven for incubations. Protease digestion was adjusted to 22 minutes and stains were developed using the ACD Dab, before being counterstained with Mayer’s Haematoxylin and Blueing Buffer. Slides were dehydrated using an ethanol series followed by Xylene and mounted using Leica Mounting Medium. High resolution 40X images were taken using the Zeiss Axio-Scan Z1.

## QUANTIFICATION AND STATISTICAL ANALYSIS

### Sequencing data processing

All raw sequencing data was converted to from bcl to fastq format using Illumina bcl2fastq software, version 2.20.0.422, with allowing up to one mismatch in each sample index barcode. Raw sequence reads were quality checked using FastQC software^9^

### Raw single cell multiomics and spatial transcriptomics data processing

For each sequenced multiomics pool, Cellranger-arc software from 10X Genomics (https://support.10xgenomics.com/single-cell-gene-expression/software/downloads/latest) was used to process, align and summarize unique molecular identifier (UMI) counts against hg38 (10x reference: refdata-gex-GRCh38-2020-A) human reference genome. On processed ATAC libraries, cellranger-arc aggr was used combined outputs from different sequencing runs into a single fragment dataset for further analysis. Matched hashing antibody panel and protein CITE-Seq (for CD4+ T-cells) data were processed together with multomics as matched feature barcoding libraries. Antibody barcodes list for Human Universal Cocktail V1 panel was downloaded from BioLegend website (https://www.biolegend.com/en-gb/products/totalseq-a-human-universal-cocktail-v1-20321).

Antibody tag UMI counts were summarized using a joint feature barcoding sequence reference of TotalSeq antibody sequences of hashing antibodies and protein expression target panel.

### Single cell multiomics data analysis

Raw UMI count, ATAC fragments and hashtag oligos (HTO) matrices were imported into R for further processing. For each multiomics sample, barcodes found in all RNA, ATAC and HTO matrices were retained. For each sample, cell calling was performed using ‘emptyDrops’^10^ function from DropletUtils to distinguish cells from empty droplets containing only ambient RNA. Furthermore, droplet barcodes for which a high percentage of total UMIs originated from mitochondrial RNAs were filtered out, as well as low total UMI count barcodes. These thresholds were derived individually for cells within each compartment following an initial clustering solution of all cells by examining and thresholding empirical distributions within each compartment, as total RNA content (notably higher in endothelial and myeloid cell populations) and mitochondrial RNA content (notably higher in epithelial cells) are highly cell type dependent.

For each individual 10x reaction, Seurat R package^11^ was used to normalize expression values for total UMI counts per cell. Highly variable genes were identified by fitting the mean-variance relationship and dimensionality reduction was performed using principal-component analysis. Cells were then clustered using Louvain algorithm for modularity optimization using kNN graph as input. Cell clusters were visualized using UMAP algorithm^12^ with principal components as input and n.neighbors = 20, spread = 1 and min.dist = 0.1. Cells from separate pools/reactions were merged together and the pool batch effect signal was corrected using the harmony algorithm^13^. Where samples from different isolation protocols were merged for further analysis (e.g., epithelial cells from EPCAM+), harmony was used to correct for 10x reaction, using equal weighting/parameters for both batch variables. In each case, merged cell clustering and visualization of cells was performed as before using Louvain and UMAP algorithms, using harmony dimensionality reduction as input instead of principal components. Merged pool clusters were compared with cell types obtained from individual pools to ensure cell type heterogeneity was not lost due to batch correction.

The ATAC fragments within BSgenome.Hsapiens.UCSC.hg38 reference genome was retained to call peaks using MACS2 CallPeaks function^14^. Signac R package^15^ was used to find the most frequently observed features, apply TFIDF to normalise ATAC peaks, reduce dimensionality using singular value decomposition (SVD), and cell clusters were visualised using UMAP algorithm. Similar to RNA, cells from different pools were merged and batch corrected using the harmony algorithm. The cell clustering and visualised was performed using harmony dimensionality reduction. The 1-50 RNA harmony dimensions and 2-30 ATAC harmony embeddings were used to construct WNN graph for RNA and ATAC joint embeddings.

The accessibility of the gene’s regulatory regions (i.e., gene score) was computed using the ArchR package^16^. The Arrow file was generated for fragments with > 4 transcription start site (TSS) enrichment score, a minimum of 1000 mapped ATAC fragments, and for cells retained after prior mitochondrial, UMI count, and ATAC filtering. The GeneScoreMatrix from the ArchR project was extracted and added to the Seurat object.

For CITE-Seq protein expression analysis, count matrices were imported into R as before as a separate assay in Seurat objects and filtered to retain only those cells passing QC based on RNA expression analysis. Hashing antibodies were split from Human Universal Cocktail antibody matrices and processed separately as described below in **Hashed sample de-multiplexing and doublet removal** section. Remaining count data were normalized using a centered log ratio transformation.

Cell populations were annotated on joint-modality clusters based on differentiation status (for epithelial cells), age-specific abundance, spatial localization (e.g. crypt top, submucosa), canonical marker genes, and previously published scRNA-seq reference atlases ^3,4,17,18^. In the case of the latter, we carried out label transfer between datasets for each cell type in reference 10x datasets for each cell in our data using Seurat and assigned putative cluster identities to populations based on the most frequently predicted cell label for each cluster. Classification results were then checked against known marker gene expression.

### Hashed sample de-multiplexing and doublet removal

Hashing antibody UMI count matrices were filtered to keep only 10x cellular barcodes from droplets passing QC based on mRNA expression profiles, as described above. Non-hashing antibody counts and hashing tags not present within any given pool (as hashed sample numbers varied between reactions between 3-5) were also filtered out for each pool individually. Each filtered matrix was used to demultiplex samples as described in^3,19^. Demultiplexed cells were visualized as tSNE plots from Euclidian distance matrixes. In each case, we then further examined whether sample demultiplexing was correct by ascertaining that the expression of sex-specific genes, such as *XIST*, segregated correctly with sample-of-origin assignments.

### MERSCOPE ST Custom Panel Design

A custom 500-gene panel was designed for MERSCOPE spatial transcriptomics to delineate over 100 fine-grained cell types within the gut, as defined by previously published single-cell reference datasets and scRNA-Seq data generated as part of this study. Genes were expert-selected to ensure comprehensive representation across stromal, immune, and epithelial lineages while excluding highly abundant transcripts to prevent optical crowding. For each identified cell type, 5 to 10 redundant genes were incorporated to ensure cell typing robustness. Probe balance was further optimised using MERSCOPE Panel Designer tool.

### MERSCOPE ST data analysis

MERSCOPE raw image data was processed and transcript decoding carried out using onboard MERLin software with custom panel codebook. For the downstream analysis of MERSCOPE *in situ* spatial transcriptomics data, we used R and the Seurat package (v5.1.0). First, raw count matrices, individual transcript coordinates, associated spatial coordinates and cell segmentations were imported into R and used to construct Seurat objects.

Next, we performed QC at cell level to filter out low-quality cells and artifacts. Cells were filtered out if they had fewer than 15 detected transcripts, if more than 10% of detected transcripts were negative control probes. Cells which were very small (< 0.01 size distribution quantile) and did not contain a nucleus, likely representing fragments, were filtered out. Cells which were very large (> 0.99 size distribution quantile) and likely represented segmentation artefacts were also filtered out. Cellular debris outside the main tissue area was filtered out by removing cells that did not have at least 5 other nearby cells. Cells in areas of other tissue artifacts, such as folds, detached areas, or areas with otherwise blurry morphology, were manually filtered out. The procedure was repeated for baysor re-segmented data (see below).

Remaining gene expression data were then normalized using Seurat’s SCTransform. For larger slides, Seurat sketch analysis was used to integrate, cluster, and project cells. Briefly, assays were split into multiple samples, and 2500 cells were sketched per sample using the SketchData function. The data were then scaled using ScaleData, and dimensionality reduction was performed with RunPCA. Integration of the data layers was achieved using the IntegrateLayers function with the RPCA method. Cells were clustered (FindClusters) and visualized using UMAP (RunUMAP). Finally, the remaining data were projected using ProjectIntegration and ProjectData to produce the final results.

### Re-segmentation of high-resolution spatial transcriptomics data

In situ spatial transcriptomics data from MERSCOPE platforms was re-segmented using Baysor^20^ algorithm, version 0.7.0. First, transcripts assigned to negative control codewords or blanks were removed. The remaining transcripts were then partitioned into cells by baysor using on-board based transcript segmentation priors. Segmentation prior confidence (--prior-segmentation-confidence) was set to 0.7. Prior cell boundary stain-based segmentation was used to calculate the average expected cell size. The number of expected clusters (--n-clusters) for the datasets was determined by Louvain clustering of original data at resolution set to 0.7.

New cell by gene counts matrices were generated from re-segmented transcript assignments. Low confidence transcripts (< 0.9) and transcripts which could not be confidently assigned to a unique cell (< 0.9) were filtered out prior to matrix generation. Gene count matrixes together with cell boundary coordinates were imported into R Seurat for further QC, clustering and visualization analysis.

### MERSCOPE ST Clustering Analysis

Cell by gene counts matrix following cell re-segmentation was converted into Seurat R object for further analysis. Very small cells, which likely represent segmentation artefacts, and cells with very few detected transcripts (<15 per cell) were filtered out from further analysis. Cells with higher than expected (>99^th^ percentile) negative probe signal were also filtered out from further analysis, resulting 1,457,779 cells in total. The full data was then dimensionality-reduced using PCA, followed by slide integration using Harmony, as described before. Nearest-neighbour graph was constructed for downstream clustering analysis and clusters were visualised using UMAP embeddings.

Low resolution clusters were annotated as broad cell type lineages based on marker gene expression and label transfer of scRNA-Seq reference data. Each broad lineage identified (e.g. Epithelium, fibroblasts, T-cells etc.) was then subset and further sub-clustered to identify cell subsets, which were similarly annotated as before. In the case where clusters were identified by our MERSCOPE panel markers that did not have a one-to-one relationship to a scRNA-Seq reference cluster and/or our panel did not contain subtype-defining markers, we annotated clusters based on their cell lineage combined with most specific gene marker expression (e.g. CXCL8+ S1/S2 and DLL+ epithelium).

### Difference Abundance Analysis

Cell-type differential abundance analysis was carried our using three different approaches. The R package sccomp (v1.9.5) was used to test cluster-level abundance differences and were visualised using ribbon and area plots using ggplot2 (v4.0.1) package. R package miloR (v2.4.1) was used to carry out graph-based differential abundance analyses based on data UMAP embeddings, an analysis which is independent of cell cluster assignments and can highlight differences between populations which may exist on a phenotypic continuum rather than falling into discrete clusters.

Alternatively, we quantified cell-type composition across samples by constructing sample-by-cluster proportion tables from single-cell and spatial data. Age groups were treated as an ordered factor spanning fetal to adult stages. To ensure robustness, we excluded sample–cluster combinations containing fewer than five cells and removed fibroblast populations derived from rectal samples. For each broad cell compartment, we calculated the relative proportion of each joint cluster within individual samples. For immune cells, as T and B cells emerge primarily during the neonatal stage, proportions were computed relative to the total immune compartment (myeloid, T, and B cells) rather than within individual immune subtypes.

To model age-associated changes in cluster proportions, we applied generalized linear models with a quasibinomial error distribution and a natural cubic spline transformation of age (4 degrees of freedom) to capture non-linear trends. Analyses were restricted to clusters represented in more than five samples and with sufficient age diversity. Predicted proportions were obtained from fitted models and aggregated across age groups. For visualization, circular heatmaps were generated using z-score–normalized values (centred and scaled across age groups) using circos.heatmap in R circlize (v0.4.16) package, with clusters annotated by their corresponding broad cell compartments.

### Differential Expression Analysis

Differential gene expression analyses, including marker gene identification, were conducted using negative binomial generalized linear models for both ST and multiomics data. To account for potential confounders, such as variability in gene detection rates and batch/donor/slide effects, these factors were incorporated as covariates into the model. Significance was assessed using Benjamini-Hochberg correction, considering genes with a false discovery rate (FDR) below 5% as significantly differentially expressed. Additionally, the MAST^21^ package was used to detect genes that exhibit more binary on/off expression patterns by modelling both continuous and discrete components.

### Pseudobulk PCA Analysis

Pseudobulk PCA analysis was carried out by aggregating UMI raw counts matrices of each individual sample by summing counts for each gene and peaks. Epithelial, immune and stromal cells were considered separately. Sample pseudobulk counts were normalised using library size factor normalisation, as implemented in DESeq2 R package (v1.50.2)^22^ for bulk RNA-Seq analysis. Counts were further transformed using variance stabilising transformation (vst) and the top 1000 most variable genes and 2000 most variable peaks were selected to compute PCA. The first two PCs were visualised to assess the overall sample-level variability within each cohort.

### Metacell construction

To address sparsity in single-cell data, we aggregated cells into metacells using the hdWGCNA^23^ framework. Genes were first filtered based on their expression within each broad cell cluster, retaining those expressed in at least 5% of cells per cluster, and genes targeted by the jump-code depletion panel were removed to prevent artefactual signals. Metacells were then generated for each sample to preserve biological heterogeneity using MetacellsByGroups on the batch-corrected RNA embedding. Ten nearest neighbors were aggregated per metacell (k = 10), allowing up to three shared cells between neighboring metacells (max_shared = 3). The metacell assay was subsequently extracted using GetMetacellObject to generate the final expression matrix for downstream analyses, including visualization of marker gene expression and building molecular clocks.

### Receptor-Ligand Analysis

Receptor–ligand interaction analysis was performed using CellChat^24^ and LIANA^25^. The analysis was conducted separately for ileal and colonic samples, and the RNA assay was filtered to retain genes with total counts greater than 50. A CellChat object was created from the filtered expression matrix to infer cell–cell communication between cell clusters. The human ligand–receptor database (CellChatDB.human) was used, restricted to secreted signalling pathways. Standard CellChat preprocessing and inference steps were applied, and communications were retained for clusters containing more than 10 cells.

LIANA analysis was performed on clusters containing more than 10 cells. Ligand–receptor communication was visualized using chord diagrams based on log fold change–weighted interaction scores. Interaction data were restricted to epithelial source cells and annotated with ligand pathway information from the CellChat human interaction database. We retained interactions belonging to selected signalling pathways, including CCL, complement, CXCL, EGF, FGF, Hedgehog, IL1, IL2, MHC-II, NOTCH, and WNT. For each age group, interactions were subset, ranked by ligand–receptor log fold change, and the top 40 interactions were visualized using chord diagrams. Chord widths were scaled according to interaction strength (log fold change), and sector colours indicated cell type. Diagrams were generated using the circlize package in R. All analyses were performed in R using the CellChat, Seurat, tidyverse, pheatmap, circlize, and RColorBrewer packages, alongside the LIANA framework implemented in Python.

### Gene Regulatory Network Analysis

SCENIC+^26^ was used to reconstruct transcription factor (TF) regulons for epithelial, immune, and stromal datasets. Cell-by-AUC matrices for each TF were imported into R for downstream analysis. The RcisTarget database containing TF motif scores for the human hg38 reference genome was downloaded from (https://resources.aertslab.org/cistarget/databases/homo_sapiens/hg38/refseq_r80/mc_v10_clust/ <u>gene_based/</u>), and the expression matrix was filtered to retain only genes present in this database.

Differential regulon activity was assessed by identifying cluster-specific regulons using FindAllMarkers. Regulons were filtered for significance (adjusted p < 0.01) and effect size (log₂FC > 0.2 and pct.1 > 0.25) to define lineage-specific regulons.

To construct a TF “decision tree” and identify transcription factors that distinguish closely related subpopulations, mean AUC values for each regulon were computed per cluster. Hierarchical clustering was performed using BuildClusterTree and hclust. Branches of the resulting cluster tree were traversed iteratively, comparing regulon specificity between child clusters at each split. Lower-level nodes were evaluated first, ensuring that a regulon assigned to a specific split was not reused at higher levels. The top regulons for each split were visualized as dendrogram edge labels. The final cluster tree was plotted as a phylogenetic tree (plot.phylo) with regulons annotated on branch edges. Regulon activity heatmaps were generated from average regulon AUCs across clusters, row-scaled, and visualized using pheatmap.

To visualise regulon activity and downstream targets, SCENIC-derived AUC assays and target gene/region memberships were extracted. TF expression and regulon activity were overlaid on UMAPs using RNA and gene-/region-level AUC assays. Downstream target genes of each regulon were tested for enrichment of Gene Ontology (GO) biological processes using clusterProfiler::enrichGO with the human annotation database org.Hs.eg.db. Results were summarized and visualized as enrichment maps (emapplot). Target gene expression and region accessibility heatmaps were generated per cluster or per age group using average RNA and peak assay matrices, scaled by row, and plotted with pheatmap. All analyses were performed in R using the packages Seurat, ape, ggplot2, igraph, ggraph, tidygraph, clusterProfiler, org.Hs.eg.db, enrichplot, and pheatmap.

### Sample-level distance to fetal anchor

All analyses were performed in Python using Scanpy for RNA preprocessing and scikit-learn and SciPy for dimensionality reduction and distance computation. To quantify how closely each sample resembled fetal tissue, we computed sample-level distances to a fetal “anchor” using RNA, ATAC (peaks), or a joint RNA + ATAC embedding. For RNA, preprocessing included library-size normalization, log₁₊ transformation, selection of the top 3,000 highly variable genes, scaling, and principal component analysis (PCA); the first 50 PCs were retained as the RNA embedding. For ATAC peaks, a term frequency–inverse document frequency (TF–IDF) transformation was applied to the sparse peak count matrix, followed by latent semantic indexing (truncated SVD); the top 50 LSI components were z-scored to produce the ATAC embedding. When both modalities were used, embeddings were computed on matched cells and concatenated.

For each sample, a centroid was defined as the mean embedding across all constituent cells, and the fetal anchor was set as the centroid of the youngest sample. Distances between each sample and the fetal anchor were then calculated using the cosine metric, yielding a pairwise sample-to-sample distance matrix.

To convert each sample’s molecular distance into a continuous “distance age”, we fitted monotonic smoothing models relating chronological age to the computed distance-to-fetal values. Distances were first linearly rescaled to a defined range ([0, 10]) using a min–max transformation. Mean distance-to-fetal values were then computed for each chronological age to reduce noise, and a monotonic isotonic regression (IsotonicRegression) was fitted to these age–distance pairs to capture the increasing trend between developmental age and molecular distance. The isotonic fit was further smoothed using a parametric nonlinear Hill function to model the sigmoidal trajectory. Fitted parameters were stored using joblib, and the final model provided a continuous encoder function mapping chronological age to scaled fetal distance.

### Cell-type maturation rate estimation

To assess age-associated maturation rates for each cell type, scaled Wasserstein distances were computed relative to a location-matched fetal reference (anchor) and modeled as a function of developmental age (log2 post-conception weeks, PCW) using loess smoothing, stratified by cell type. Smoothed trajectories were fit using a fixed span parameter (0.75). For each cell type, peaks were defined as the maxima of the fitted loess curves, while inflection points were identified from zero-crossings of the numerically estimated second derivative. Visualizations included individual observations, loess-fitted trajectories, and annotated peak and inflection coordinates, with developmental age intervals indicated by shaded background regions corresponding to age groups. All analyses and visualizations were performed in R using tidyverse, ggplot2, viridis, ggpubr, and related packages.

### Molecular maturation clock – ElasticNet model

Leveraging single-cell multi-omics data that capture coordinated transcriptional and epigenetic changes from fetal to adult stages across epithelial, stromal, and immune compartments, we sought to build a single-cell molecular clock to predict gut maturation and to identify genetic and epigenetic features that are predictive of this process. To this end, we adopted two complementary strategies: (i) training a multimodal Elastic Net model on our multi-omics dataset to highlight epigenetic regulations, and (ii) training a deep learning–based multilayer perceptron (MLP) regressor on a mega-transcriptomic developmental atlas with enhanced coverage of early developmental stages to improve robustness and predictive accuracy.

To construct a multimodal ElasticNet model, we first mitigated single-cell sparsity—particularly in ATAC data—by aggregating transcriptionally similar cells from the same donor into metacells. We first subset cells to the tissue of interest (either colon or ileum). To avoid sex-chromosome confounding, genes matching XIST/TSIX or a curated Y-chromosome genes were removed from all layers. A set of canonical B-cell/ plasma markers (immunoglobulin chains, JCHAIN, MZB1, PRDM1, IRF4, TNFRSF17, POU2AF1, CD38, etc.) were also de-souped to avoid confounding effect of highly expressing immunoglobulin and plasma genes. Metacell objects were generated from RNA, peaks and regulon matrices. The metacell counts were normalized, log transformed. The distance encoder was used to convert age into distance-to-fetal age, to better reflect the data-driven distance. An age-matched held-out test set was pre-defined and used to split processed AnnData objects into training and tests subsets for subsequent model training and evaluation.

For each assay (RNA, ATAC, and regulon activity), features were derived using standardized dimensionality reduction workflows. ATAC matrices were transformed by term frequency–inverse document frequency (TF–IDF), reduced by truncated singular value decomposition (LSI), and L2-scaled. RNA features were obtained by TF–IDF + SVD. Regulon activities were scaled and, when appropriate, reduced by PCA or SVD. A supervised ElasticNet regressor was trained to predict developmental age from these features. In the cell-level pipeline, cells were randomly partitioned into training and test sets (default test fraction = 0.3). The model was trained on cell-level features, and predictions on held-out cells were evaluated by mean absolute error (MAE) and other calibration metrics. In the donor-level pipeline, data were split by donor before training. All preprocessing and dimensionality-reduction steps were fitted on training donors and applied to test donors only. Cell-level predictions were averaged per donor to obtain donor-level predicted and true ages, from which donor-level MAE and summary metrics were computed.

We also implemented a two-stage regression framework to predict developmental age by integrating RNA and chromatin or regulon features. In Stage A, a baseline ElasticNet model was trained on RNA-derived embeddings alone. RNA features were computed by TF–IDF + SVD. The model was trained on all RNA cells from the training set and evaluated on held-out test cells; mean absolute error (MAE) was recorded as the baseline RNA performance. In Stage B, cells profiled with both RNA and either ATAC peaks or regulon activities were used to refine predictions through feature fusion. ATAC features were derived by TF–IDF followed by latent semantic indexing (truncated SVD), and regulon activities were scaled and optionally reduced by PCA. The pretrained RNA embeddings were aligned with the corresponding ATAC or regulon features for the same cells, and fusion was performed using residual learning. The residual learning modeled the remaining error from the RNA-only predictor (the “residual”) as a function of additional chromatin or regulon features. The RNA model provided a base prediction for each cell, and a second ElasticNet model learned to correct systematic deviations by fitting the residuals against the secondary modality. The corrected prediction was then obtained by summing the RNA base prediction and the learned residual term, effectively capturing modality-specific variance not explained by RNA alone. All preprocessing and transformations were fitted on the training set and applied unchanged to the test set. Performance was assessed at both the cell and donor levels, with donor-level predictions obtained by averaging per-cell outputs across donors. All analyses were implemented in Python.

### Molecular maturation clock – MLP Regressor

First, to extend developmental representation, we integrated single-cell RNA-seq datasets from Elmentaite et al.^27,28^ and Fawkner-Corbett et al.^2^, excluding extreme preterm samples, yielding 41 colonic and 59 ileal samples for training and validation. We then trained a deep learning–based multilayer perceptron (MLP) regressor on a mega-transcriptomic developmental atlas.

We trained a multilayer perceptron (MLP) regression model to predict developmental age from gene expression profiles. AnnData objects were filtered to retain cells expressing at least 200 genes and genes detected in at least three cells, followed by library-size normalization to 10,000 counts per cell and log1p transformation. The training and test datasets were merged for shared preprocessing, and the top 5,000 highly variable genes were selected. Expression values were then z-score scaled. To support downstream visualization and batch correction, principal component analysis was performed, and batch effects were harmonized using Harmony; nearest-neighbor graph construction, UMAP embedding, and Leiden clustering were subsequently applied.

The age-prediction model was trained on the processed training set using an MLP with three hidden layers (512, 256, and 128 units), batch normalization, ReLU activation, and dropout (0.2). The model was optimized with AdamW using an L1 loss function, a learning rate of 1 × 10^-3, and a batch size of 64. Data were split into training, validation, and test sets, and early stopping was applied based on validation mean absolute error (MAE) with a patience of eight epochs. Performance was evaluated using MAE and R^2 on held-out test data. For application to the independent test dataset, genes were aligned to the training feature space, missing genes were imputed as zeros, and expression values were scaled using the training mean and standard deviation before prediction. Predictions were also aggregated at the sample level using both mean and median summaries.

To interpret model predictions, integrated gradients were computed using a zero-expression baseline and 100 interpolation steps. Gene-level attribution scores were summarized across cells to generate signed and absolute importance metrics, and top-ranked genes were visualized. Single-cell preprocessing was performed using Scanpy^29^ and AnnData, with batch correction using Harmony^13^. Neural network modelling was implemented in PyTorch^30^. Model evaluation, numerical analysis, visualization, and data handling were carried out using scikit-learn^31^, SciPy, NumPy, Pandas, Matplotlib, Seaborn, and tqdm.

### Pathway Enrichment Analysis

Gene Ontology (GO) and Reactome pathway enrichment analyses for both scRNA-Seq and spatial transcriptomics data were conducted using the clusterProfiler R package^32^ (v4.18.4). Gene identifiers were mapped with the org.Hs.eg.db (v3.22.0) annotation DBI package. Individual cluster marker sets and differentially expressed genes were analysed for overrepresentation, using all expressed or detected genes as the background reference. Hypergeometric p-values were adjusted for multiple testing through the Benjamini–Hochberg method. The enrichment results were visualized with clusterProfiler and ggplot2 packages. For pathway activity analysis at the single-cell or spot level, pathway scores were derived as gene module scores using the AddModuleScore function in the Seurat R package.

### Pathway Activity Scoring

Reactome gene sets were retrieved from MSigDB (category C2, subcategory CP:REACTOME) and pre-filtered to remove small gene sets containing fewer than 20 genes. To reduce redundancy, gene sets that were strict subsets of larger gene sets were identified and excluded. Long pathway names were abbreviated where appropriate. Pathway module scores were computed on the RNA assay of ileal epithelial cells for each curated Reactome gene set, using a threshold of 0.2 to define cells with elevated pathway activity.

Age-dependent pathway activity was visualized using network graphs. Pairwise pathway similarity was calculated as the proportion of shared genes between gene sets, and edges with similarity greater than 0.05 were retained. The resulting similarity matrix was converted into an undirected weighted graph, and Louvain clustering was applied to identify pathway communities. Node size was scaled to pathway gene set size. For each pathway, the fraction of cells in each age group with a module score greater than 0.2 was calculated, normalized to sum to one, and represented as pie charts at graph nodes. Network plots were generated using ggnetwork with geom_scatterpie (node radius scaled as log₂(node_size) × 0.0025), where edge width reflected gene overlap and node colour indicated the age group with elevated pathway activity. All analyses were performed in R using the packages msigdbr, Seurat, igraph, ggplot2, and pheatmap.

### Consensus Non-negative Matrix Factorisation (cNMF)

To identify transcriptional programs in epithelium, consensus non-negative matrix factorization (cNMF)^33^ was applied to gene expression matrices. Prior to factorization, Harmony was used to correct for sequencing-specific batch effects. cNMF was performed to extract 15 factors, each representing a distinct gene expression program within the epithelial dataset. The resulting gene loadings and usage scores were associated with known variables, including diet, sex, age, location and cell types. To quantify the contribution of biological factors to variation in gene program activity, we applied variance partitioning using the variancePartition R package. Normalized program usage scores were transposed into a cell-by-program matrix, and a linear mixed model was fitted including sex, age group, diet, and location. For each program, the model estimated the proportion of variance attributable to each factor. The resulting variance fractions were visualized using plotVarPart, and factors were ranked by their mean variance explained across programs.

Associations between module scores (X1–X15) and metadata were assessed using Pearson correlation. Age and scaled pseudotime were treated as continuous variables, whereas diet, cell type and location (Colon, Ileum) were one-hot encoded; correlations were computed using pairwise complete observations and visualized as hierarchically clustered heatmaps (pheatmap).

### Epithelial Differentiation by Pseudotime Analysis

The marked separation of fetal enterocytes prompted us to ask whether these cells represent immature intermediates or a developmentally distinct mature population restricted to early gestation.

To disentangle differentiation state from developmental time, we modelled enterocyte maturation along two orthogonal axes: crypt-axis differentiation and chronological age. As age-associated transcriptional differences plateaued at approximately 8 months in the ileum, and epithelial profiles from fetal, <6-month, and >6-month samples clustered tightly within each group in principal component space, we reconstructed pseudotime trajectories separately for these age groups.

Pseudotime trajectories were constructed for six groups spanning intestinal region (colon and ileum) and developmental stage—fetal, infants (<6 months), and paediatric (>6 months). For each group, Seurat objects were pre-processed, batch-corrected, and integrated to generate low-dimensional embeddings. Pseudotime trajectories were then computed using the Monocle 3 algorithm (Qiu et al., 2017), with stem cell populations defined as the root nodes to capture differentiation dynamics along absorptive and secretory epithelial lineages. Pseudotime values from all age groups were subsequently scaled, merged, and projected back onto the original UMAP embeddings for both the ileum and colon, providing a unified view of epithelial differentiation across development.

### Cross-modal Embedding Distance

To quantify the concordance between transcriptional and chromatin accessibility structures, we calculated per-cell Wasserstein distances between RNA- and ATAC-derived neighbourhoods. Batch-corrected embeddings were extracted from the RNA and Peak assays, and the first 30 dimensions of each were retained. For every cell, k-nearest neighbours (k = 200) were identified independently in the RNA and ATAC spaces (excluding the self-neighbour), and the resulting neighbour-distance vectors were compared. The one-dimensional Wasserstein distance (wasserstein1d) between each cell’s RNA and ATAC distance distributions was computed and stored as a per-cell concordance score for downstream analysis.

To relate chromatin–transcriptional concordance to epithelial state, cells were stratified by marker expression: MKI67 ≥ 0.5 (MKI67), LGR5 ≥ 0.5 and MKI67 ≥ 0.5 (LGR5⁺MKI67⁺), and LGR5 ≥ 0.5 with MKI67 = 0 (LGR5⁺MKI67⁻). Remaining cells were categorized as differentiated or undifferentiated according to pseudotime position relative to the stem node (see **Epithelial differentiation by pseudotime analysis**). Wasserstein distance distributions across these epithelial states were visualized as violin plots in Seurat. All analyses were conducted in R using the packages FNN, transport, Seurat, RANN, and ggplot2.

### TF In silico Perturbation

Gene regulatory network (GRN) inference and in silico perturbation analyses were performed using CellOracle^34^ in epithelial cells. Raw count matrices were imported into a CellOracle object together with UMAP embeddings and cell-type annotations, and a prior GRN was provided using a binary eRegulon matrix. After principal component analysis, gene expression was denoised by k-nearest-neighbor imputation using an automatically selected number of components and neighborhood size. Cell-type–specific GRNs were then inferred and filtered based on statistical significance (p < 0.001) and edge weight.

To model developmental trajectories, pseudotime values derived from Monocle3 were integrated and used to construct a developmental vector field. In silico perturbation simulations were performed by setting expression of selected transcription factors (e.g. TCF7L1 and ARID3A) to zero and propagating their effects through the inferred GRNs. Simulated cell-state transitions were compared to the developmental vector field, and perturbation effects were quantified using inner-product scores across the embedding space. Analyses were performed in Python using CellOracle.

### Annotation of T-cell subsets, states and functions

StarCAT^35^ is a reference-based annotation framework that quantifies predefined gene expression programs (GEPs) to characterize cellular identity and functional states from single-cell transcriptomic data. It enables annotation of T cell subsets, activation states, and functional programs by scoring cells against curated signatures derived from large-scale reference datasets. StarCAT analysis was performed using the web-based implementation on T cell and intraepithelial lymphocyte (IEL) populations. Default parameters were used.

### Promoter and enhancer accessibility

Promoter and enhancer genomic regions were defined using hg38 promoter annotations and GeneHancer enhancer–gene interaction databases^36^, respectively. Promoter regions were defined as ±1.25 kb around transcription start sites, and chromatin accessibility across these regions was quantified from single-cell ATAC-seq data by intersecting annotated intervals with ATAC-seq peaks. Accessibility profiles were visualized using coverage plots and modeled as a function of age using generalized additive models (GAMs).

To relate differential gene expression to regulatory accessibility, we analyzed promoter and enhancer accessibility of genes exhibiting heightened IFNγ or TNFα responses in organoids. The same genomic annotations were used, and duplicate enhancer entries were collapsed prior to analysis. Accessibility profiles were extracted from Seurat CoveragePlot tracks and summarized across age groups using the area under the coverage curve (AUC), computed by trapezoidal integration, together with summary statistics of coverage. Age-associated changes in accessibility were modeled for each gene–region pair using linear regression and GAMs (spline basis, k = 4), and regions with low accessibility signal were excluded. Genes were classified as increasing or decreasing when the GAM term was significant and the linear slope was positive or negative, respectively, and as no trend otherwise; these trends were then compared with corresponding gene expression patterns. All analyses were performed in R using Seurat, Signac, and associated packages.

### ST Crypt Niche Detection

Cell niche detection was performed using an unsupervised approach implemented in the ‘spamsc’ R package (https://github.com/ChloeHJ/spamsc). For each field of view in the spatial transcriptomics dataset, spatial coordinates of cell centroids were extracted to determine the 30 nearest neighbours of each cell. Using these coordinates, two neighbourhood matrices were generated: (1) expression counts from each cell’s nearest neighbours were aggregated to produce a cell-by-gene matrix representing the local transcriptomic environment, and (2) the cell types of neighbouring cells—defined by high-resolution subcluster annotations from previous analyses—were counted and aggregated into a cell-by-cell-type matrix representing the local cellular composition. This process was repeated for all slides, and resulting matrices were merged across fields of view.

To identify spatial niches, unsupervised clustering was performed on the neighbourhood matrices. Principal component analysis (PCA) was applied to the gene expression matrices, followed by batch correction and integration of principal components across slides using Harmony. Niche clusters were then detected using Louvain community clustering. Crypt niches were defined as those predominantly composed of epithelial cells.

### ST Cell Co-localisation Analysis

Spatial niche analysis was performed on MERSCOPE ST data as previously described. Cellular niches were first defined by aggregating gene expression from the 30 nearest neighbouring cells (k = 30) using spatial coordinates, followed by clustering of niche expression profiles.

To characterize local neighbourhood relationships within the crypt niche, spatial co-localisation between cell subclusters was quantified using precomputed nearest-neighbour pairs. For each reference cell, neighbouring cell subclusters were aggregated across all cells, and counts were converted into proportions within each reference subcluster, generating a row-normalized adjacency matrix describing the frequency of spatial associations.

Interaction matrices were filtered to retain cell subclusters with a minimum total interaction proportion of 0.2 and edges with a minimum weight threshold of 0.1. The resulting matrices were represented as undirected weighted networks, where nodes correspond to cell subclusters, node size reflects cell abundance, and edge thickness represents the strength of co-localisation. Nodes were coloured by cell compartment, and networks were visualized using force-directed layouts. Analyses were stratified by developmental age groups (fetal, <1 month, <2 months, and >2 months) to assess temporal changes in spatial organization. All analyses were performed in R using Seurat, tidyverse, igraph, ggraph, and ggforce.

### ST adjacency-based epithelial–neighbourhood correlation analysis

To assess interactions between epithelial cells and their local microenvironment, we quantified correlations between epithelial gene expression programs and neighbouring cell–derived signals across samples. Age groups were consolidated into three categories (fetal, <1 month, and >1 month), and analyses were restricted to epithelial cells from the terminal ileum, with selected outlier samples excluded. For each sample, expression profiles of neighbouring cells (cytokine-related genes) and epithelial cells (genes representing pathways of interest) were extracted, and Pearson correlation matrices were computed between neighbour-derived signals and epithelial gene expression.

To obtain robust estimates, correlation matrices were averaged across samples within each age group, and corresponding standard deviations and standard errors were calculated. Age-specific average correlation matrices were visualized as heatmaps with hierarchical clustering, using a consistent global colour scale across age groups.

### Bulk RNA-seq analysis

Raw sequence reads were quality checked using FastQC software^9^. Cutadapt^37^ was used to trim poor-quality bases and Illumina universal adapter sequences from raw reads before alignment.

The human hg38 reference genome analysis set was obtained from the University of California Santa Cruz (UCSC) ftp site^38^. For BHLHE40 experiment, full lentiviral construct sequences (Addgene plasmids #142336) were included as additional reference contigs and indexed together with hg38 as a reference genome using STAR aligner^39^. Reads were then aligned to this custom reference.

Raw gene expression counts were summarized with featureCounts^40^ using a custom hg38 and plasmid GTF file containing joint annotations. The MultiQC^41^ tool was used to aggregate quality metrics. Sample quality metrics and raw read counts were imported into R for further processing. The DESeq2 R package^22^ was used to estimate library size factors, normalize counts and perform differential expression analyses. Benjamini–Hochberg multiple testing correction was used to compute FDR, and genes were considered significantly differentially expressed at <5% FDR. PCA was performed in R using the top 1,000 most variable genes, with normalized DESeq2 variance-stabilized transformation expression as input. Combat [85] was used to correct donor-specific effects for heat map visualizations only.

## Supplementary Figure Legends

**Supplementary Figure 1.**
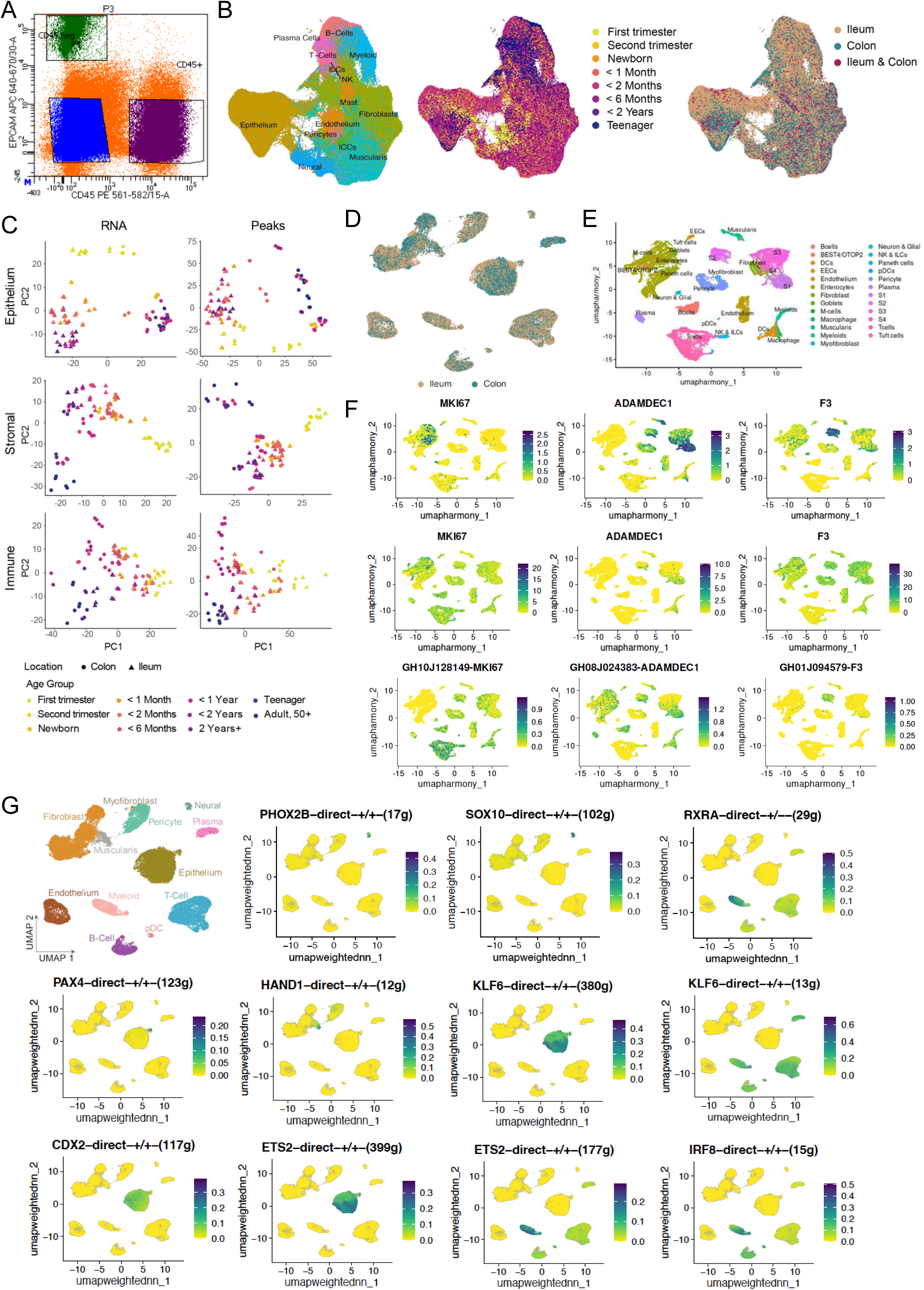
Multimodal characterisation using single cell multiomics and ST data. A. Flow cytometry gating strategy for isolation of stromal (CD45⁻) and immune (CD45⁺) compartments. B. UMAP of integrated MERSCOPE ST data, coloured by broad cell type, age group and location. C. Pseudobulk principal component analysis (PCA) of RNA and peak assays across epithelial, stromal and immune compartments identified by single cell multiomics, coloured by age group. D. UMAP of integrated single cell multiomics data, coloured by location. Paired UMAP overlay with cell cluster and age is shown in Figure 1C. E. UMAP of metacells derived from integrated single cell multiomics data, coloured by cell type. F. UMAP overlays showing gene expression (top), promoter accessibility (middle) and enhancer accessibility (bottom) for marker genes (MKI67, ADAMDEC1, F3). G. UMAP overlay showing cell type annotations and regulon activity in scDOGMA-seq data. Transcription factor regulons mark distinct lineages, including PHOX2B and SOX10 (enteric nervous system), HAND1 (mesenchymal), IRF8 (immune), PAX4 (endocrine) and CDX2 (epithelial), whereas KLF6 and ETS2 show broad activity across cell types.

**Supplementary Figure 2.**
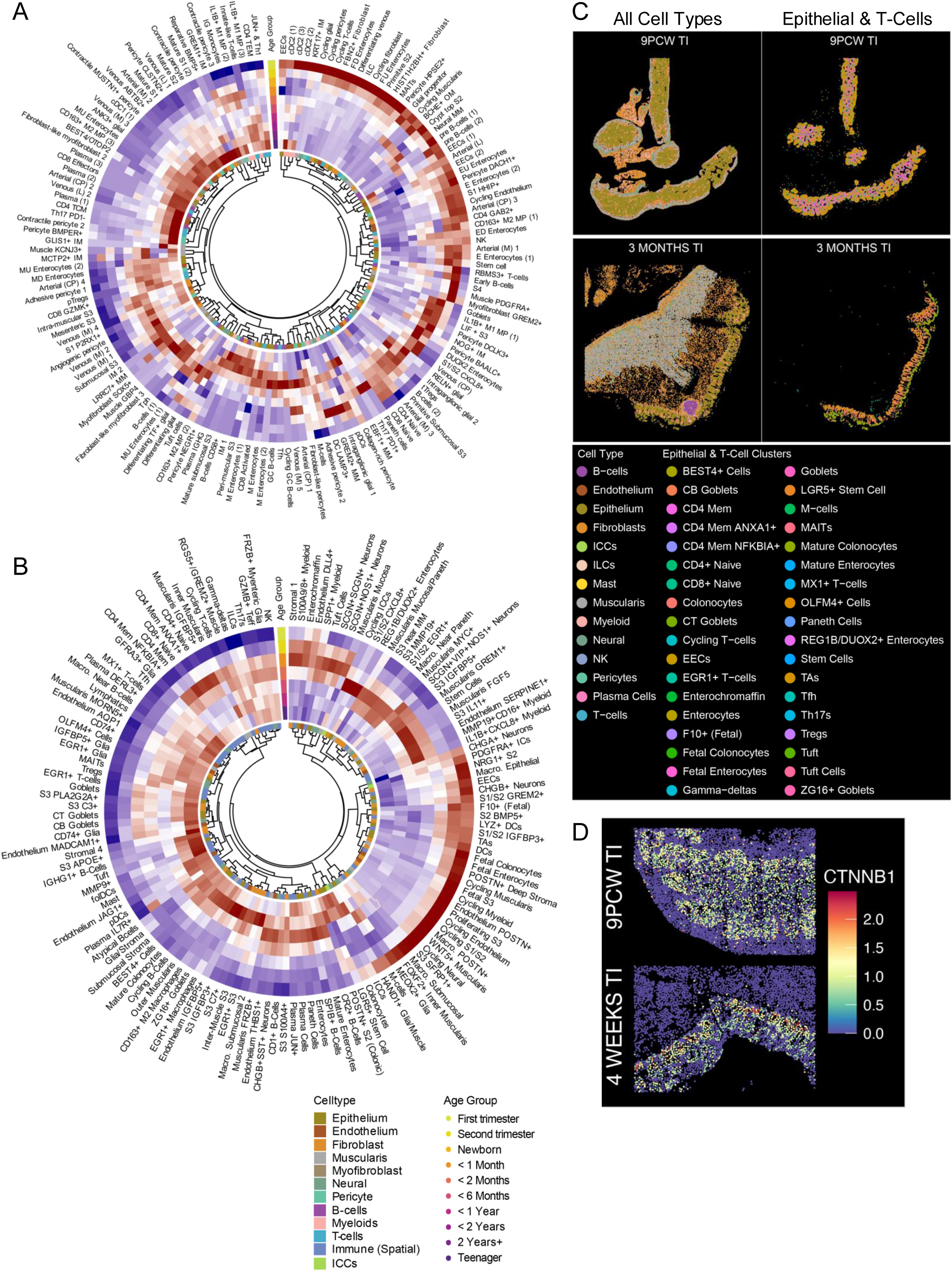
Age-associated changes in cell-type composition in single cell multiomics and ST data. A. Circular heatmap of predicted cell-type proportions from single cell multiomics. Age effects were modelled using quasibinomial generalized linear models with a natural cubic spline for age; predicted proportions are z-score normalized across age groups and annotated by broad compartments. B. Circular heatmap of predicted cell-type proportions from MERSCOPE ST data, z-score normalized across age groups and annotated by broad compartments. C. Representative MERSCOPE ST sections from 9 PCW (top) and 3 months (bottom) terminal ileum, coloured by broad cell type (left) and epithelial and T cell clusters (right). D. Spatial expression of CTNNB1 in MERSCOPE ST sections at 9 PCW and 4 weeks.

**Supplementary Figure 3.**
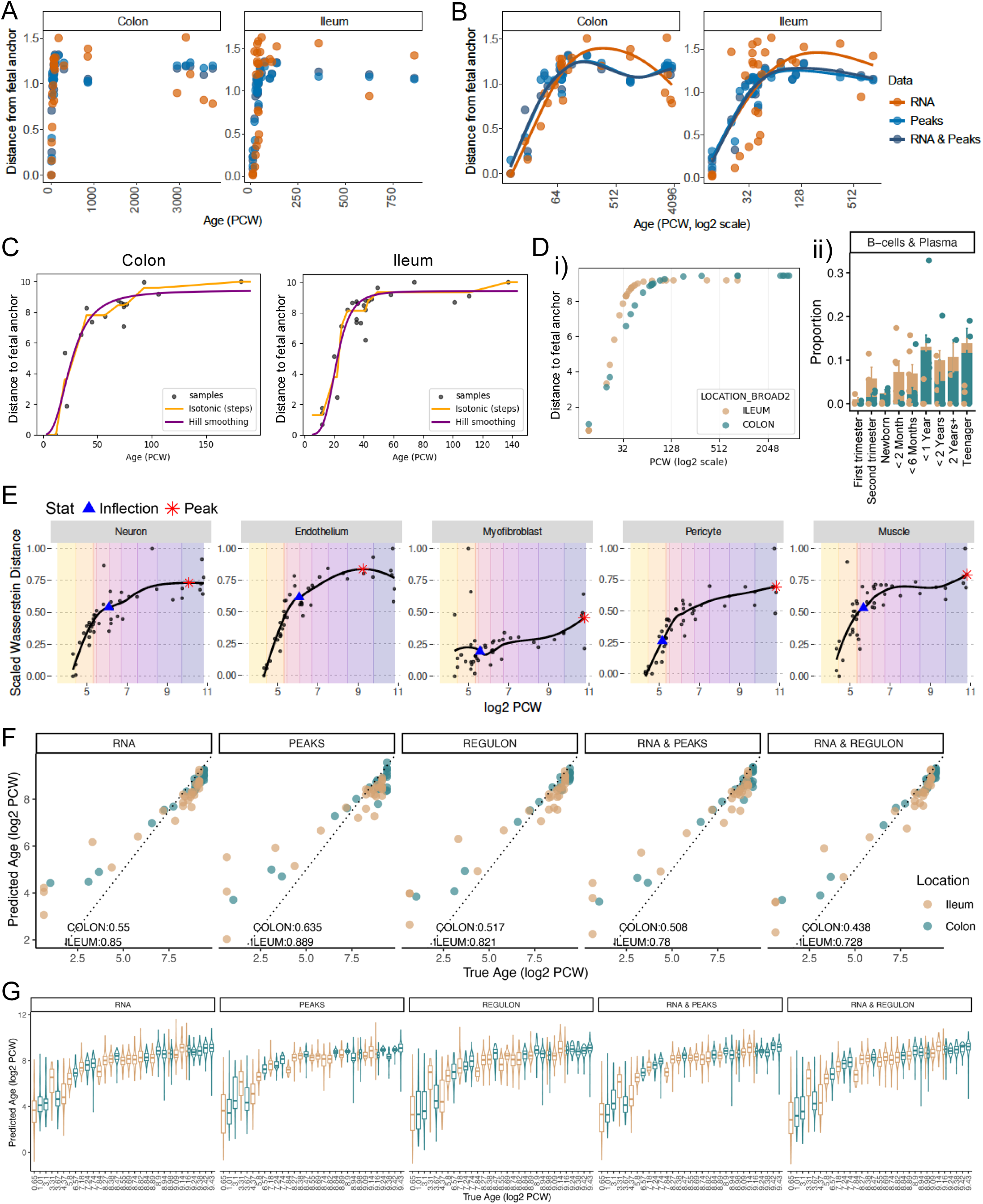
Modelling developmental trajectories and maturation rates. A. Cosine distance to location-matched fetal anchors for each donor, computed from gene expression (RNA), chromatin accessibility (ATAC peaks), and concatenated RNA+ATAC embeddings. Samples are plotted by developmental age (post-conception weeks, PCW), with each point representing a donor-level centroid. Fetal anchors were defined from the youngest samples within each tissue, and distances reflect divergence from the fetal reference state. B. As in A, plotted against log2-transformed PCW with generalized additive model (GAM) fits. C. Isotonic regression with Hill smoothing; each point represents a sample. D. (i) Location-specific maturation rates inferred from Hill-smoothed fits. (ii) Proportions of B cells and plasma cells within CD45+ compartment in colon and ileum. E. Scaled Wasserstein distance from a location-matched fetal anchor plotted against log2-transformed post-conception weeks (PCW). Each point represents an individual donor. LOESS fits are shown, highlighting the inflection and peak of the trajectories. F. Performance of a location-matched ElasticNet model comparing median predicted and true age per donor (log2 PCW), coloured by location. G. Distribution of ElasticNet performance across locations; box plots indicate quartiles (Q1, median, Q3).

**Supplementary Figure 4.**
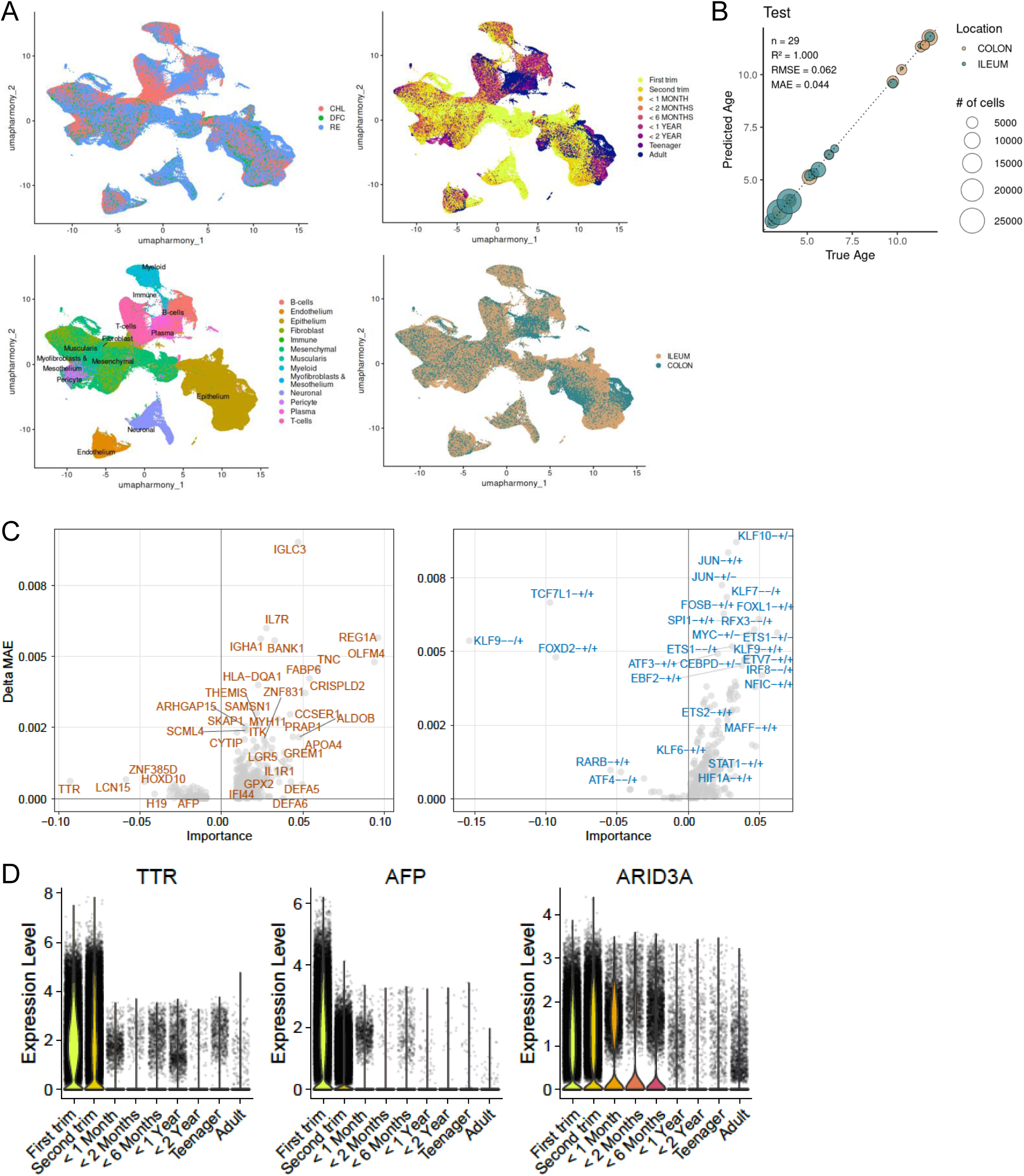
Cross-cohort integration and molecular clock performance. A. UMAP of integrated single-cell transcriptomic data with datasets from Elmentaite et al. and Fawkner-Corbett et al., coloured by data source, age group, broad cell type and location. B. Performance of a multilayer perceptron (MLP) regressor on a test dataset. Predicted age corresponds to the median across cells per donor. Points are coloured by location and sized by cell number. C. Volcano plots showing top genes (left) and regulons (right) from the ileal ElasticNet model. Axes indicate signed feature importance and change in mean absolute error (MAE) following in silico perturbation. D. Violin plots of selected fetal marker gene expression across age groups.

**Supplementary Figure 5.**
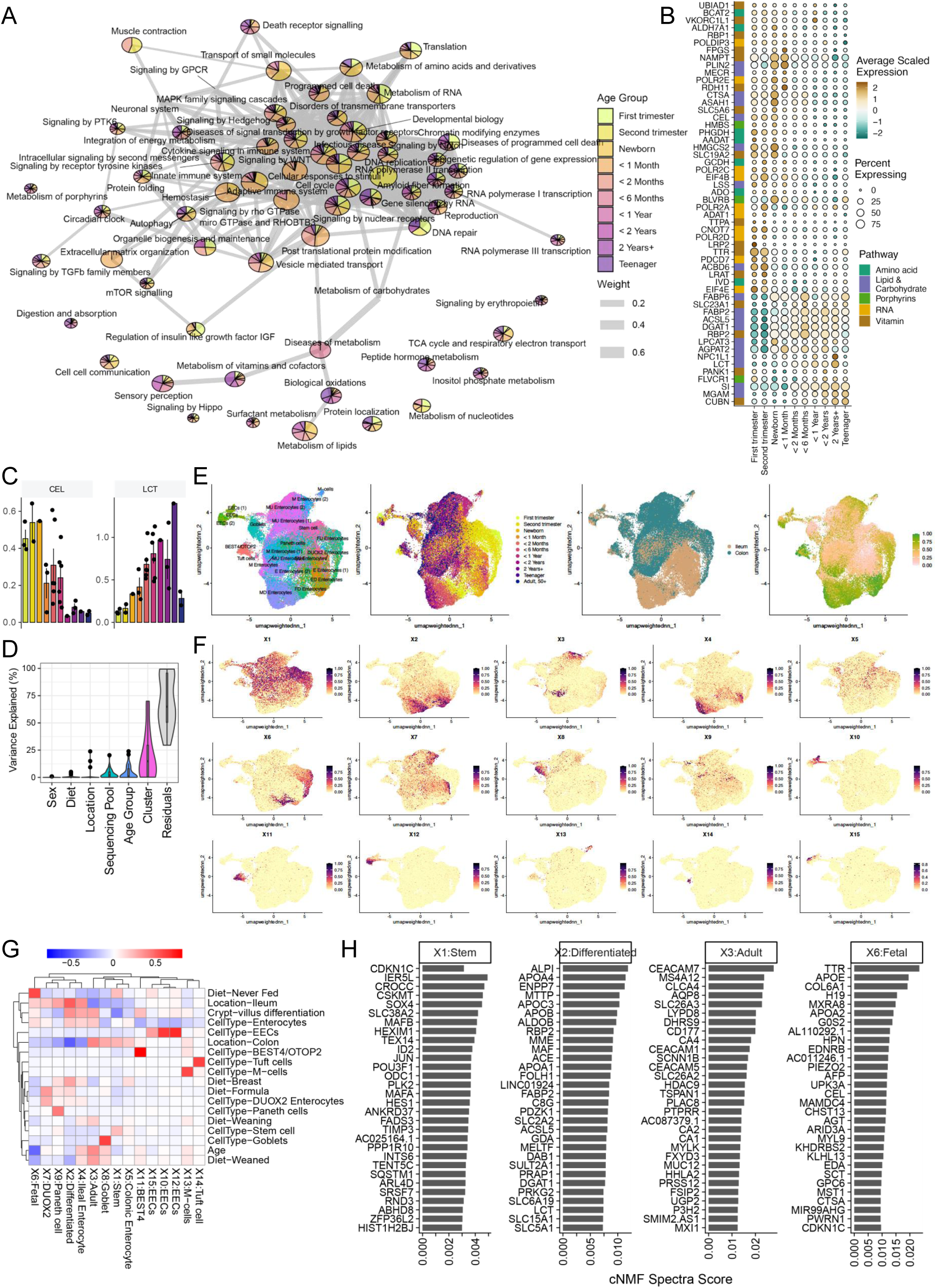
Metabolic programmes and module associations across development. A. Network representation of age-dependent pathway activity. Pathway similarity was defined as the proportion of shared genes between gene sets, retaining edges with similarity > 0.05. The network was clustered using the Louvain algorithm. Node size reflects gene set size, edge width indicates gene overlap, pie charts show the normalized fraction of cells in each age group with module scores > 0.2, and node colour denotes the age group with highest pathway activity. B. Bubble plot showing expression of metabolic pathway genes across age groups. C. Bar plot showing mean expression of selected metabolic genes across donors within each age group; points represent individual donors. D. Violin plot showing variance explained by biological and technical covariates. E. UMAP of epithelial cells from colon and ileum, coloured by cell type, age group, location and crypt–villus differentiation pseudotime. F. UMAP showing cNMF programme usage. G. Heatmap of Pearson correlations between module scores (X1–X15) and metadata features, including age, crypt–villus differentiation, diet, cell type and location. H. Bar plot showing the top 30 marker features of selected cNMF programmes.

**Supplementary Figure 6.**
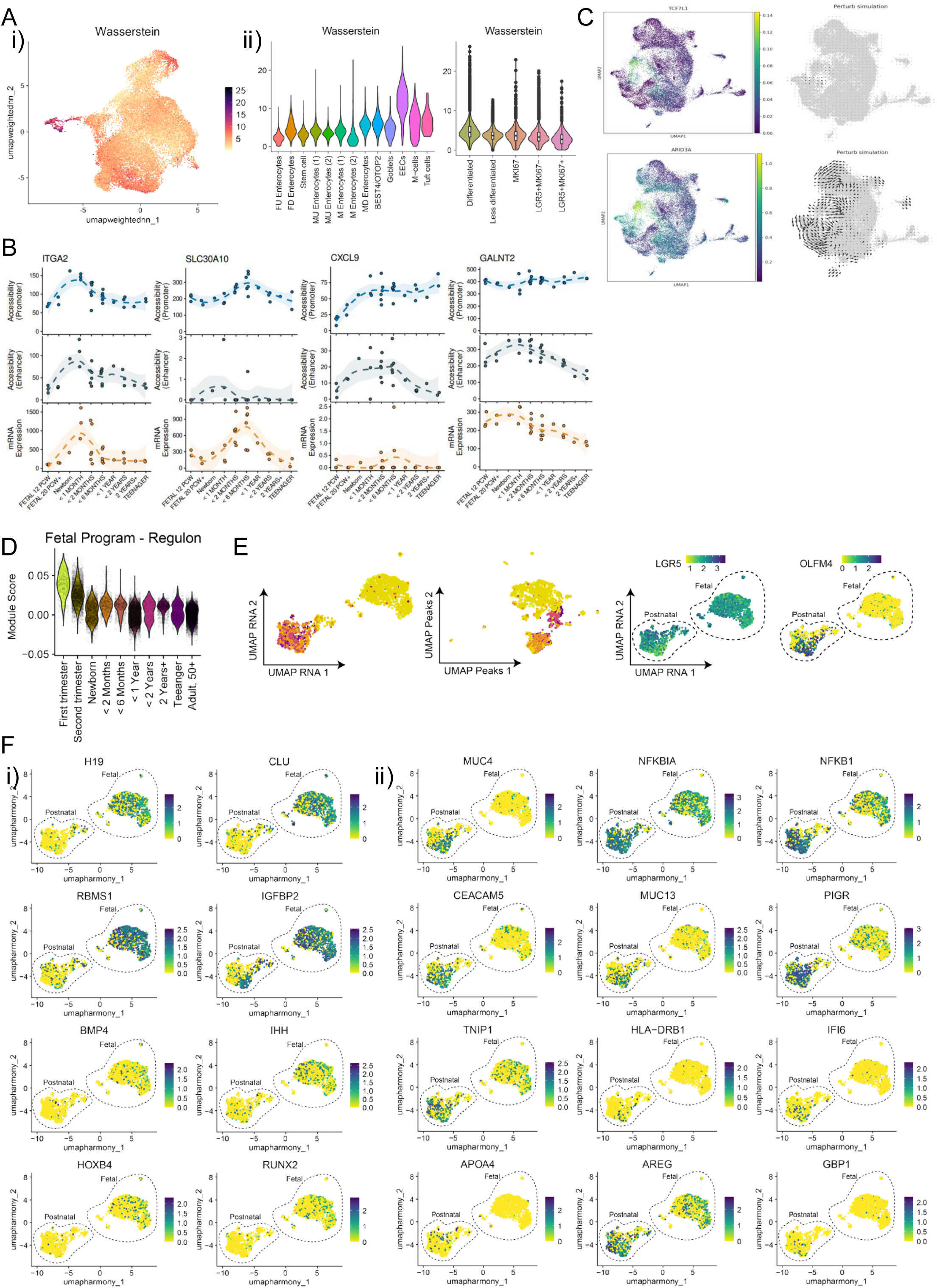
Fetal regulatory programs and stem cell dynamics across development. A. RNA–ATAC concordance assessed by Wasserstein distance. (i) UMAP of colonic epithelial cells coloured by RNA–ATAC Wasserstein distance. (ii) Distribution of distances across cell types. B. GAM-fitted trends showing promoter and enhancer accessibility per sample and gene expression of selected genes across age in ileal epithelium. Shaded areas indicate +/- 1 standard error of the GAM fitted mean. C. UMAP showing expression of fetal transcription factors (TCF7L1, ARID3A) and simulated cell-state transitions following in silico perturbation (CellOracle). D. Violin plots showing fetal regulon activity module scores in colonic epithelium. E. UMAP of colonic stem cells in RNA and ATAC embeddings, coloured by age group. (ii) UMAP showing LGR5 and OLFM4 expression in RNA embeddings. F. UMAPs showing expression of selected fetal (i) and paediatric (ii) stem cell marker genes.

**Supplementary Figure 7.**
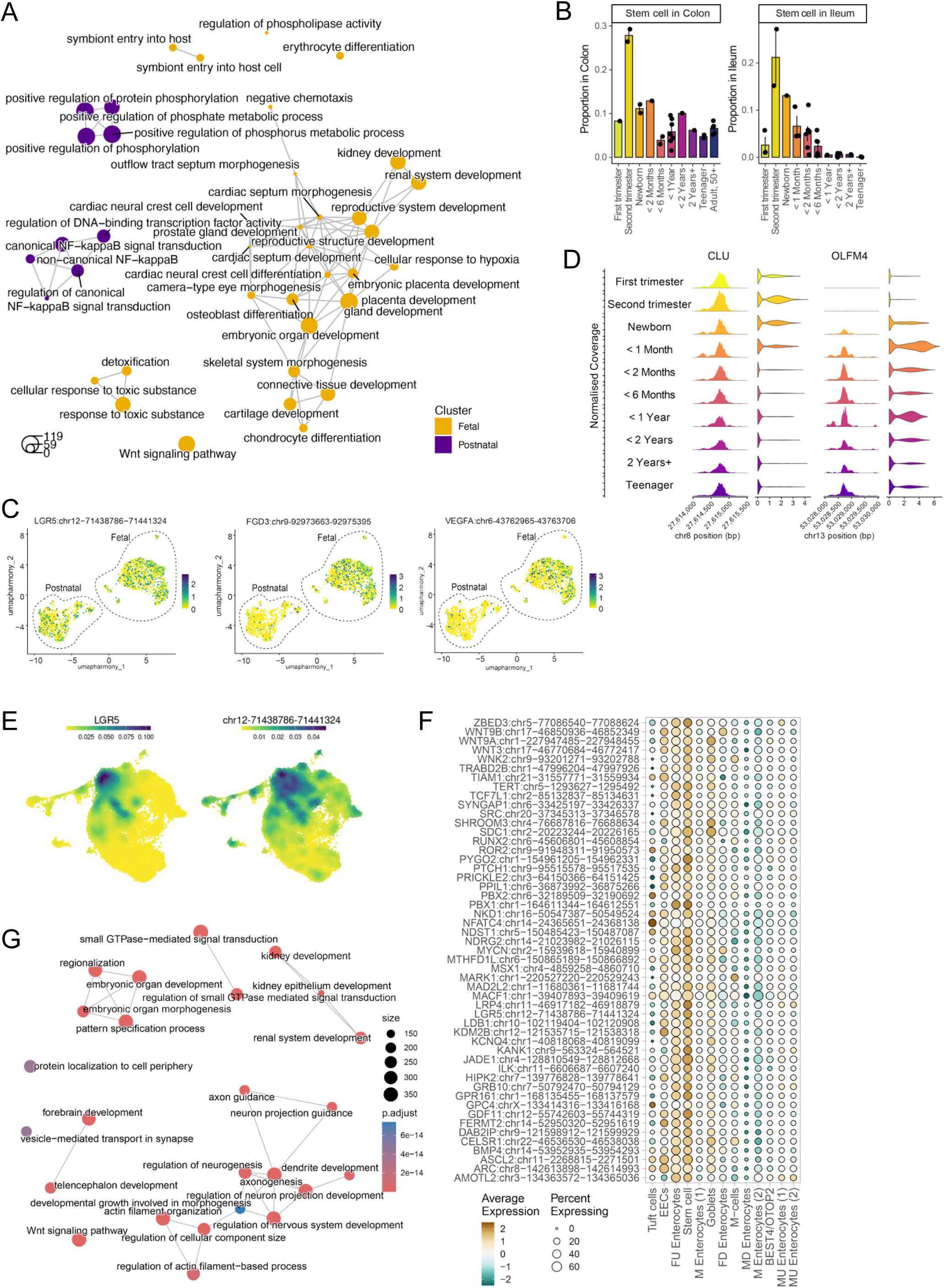
Fetal and paediatric stem cell programmes. A. Differential pathway network comparing fetal and paediatric colonic stem cells. B. Proportion of stem cells in colonic and ileal epithelium across age groups; points represent donors and bars indicate mean proportions. C. UMAP showing LGR5 promoter accessibility and selected fetal stem cell marker peaks. D. Coverage and violin plots showing chromatin accessibility and gene expression of fetal and paediatric stem cell marker genes. E. Density plots of LGR5 gene expression (left) and promoter accessibility (right) in ileal epithelial cells. F. Bubble plot showing accessibility of stem cell marker peaks. G. Reactome pathway network of genes linked to stem cell marker peaks.

**Supplementary Figure 8.**
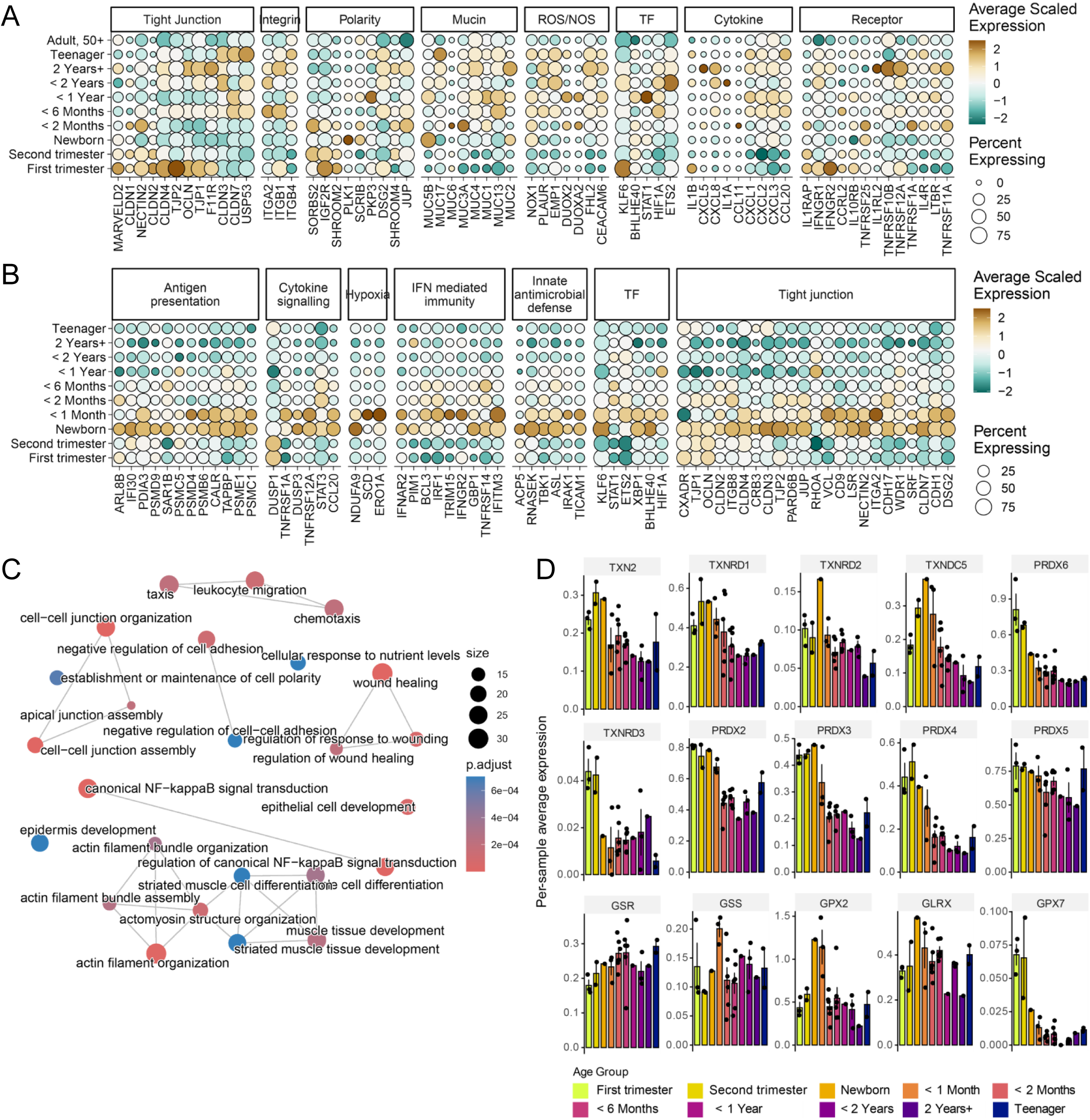
Epithelial gene expression and regulatory programmes. A. Bubble plot showing expression of genes associated with tight junctions, integrins, polarity, mucins, reactive oxygen species (ROS), nitric oxide (NO)-related pathways, transcription factors, cytokines and receptors in colonic epithelial cells. B. Bubble plot showing expression of genes included in the transcription factor regulatory network (Fig. 3B) in ileal epithelial cells. C. Network representation of pathways associated with KLF6 downstream target genes. D. Bar plot showing mean expression of selected thioredoxin and glutathione pathway genes in ileal epithelial cells across donors within each age group; points represent individual donors.

**Supplementary Figure 9.**
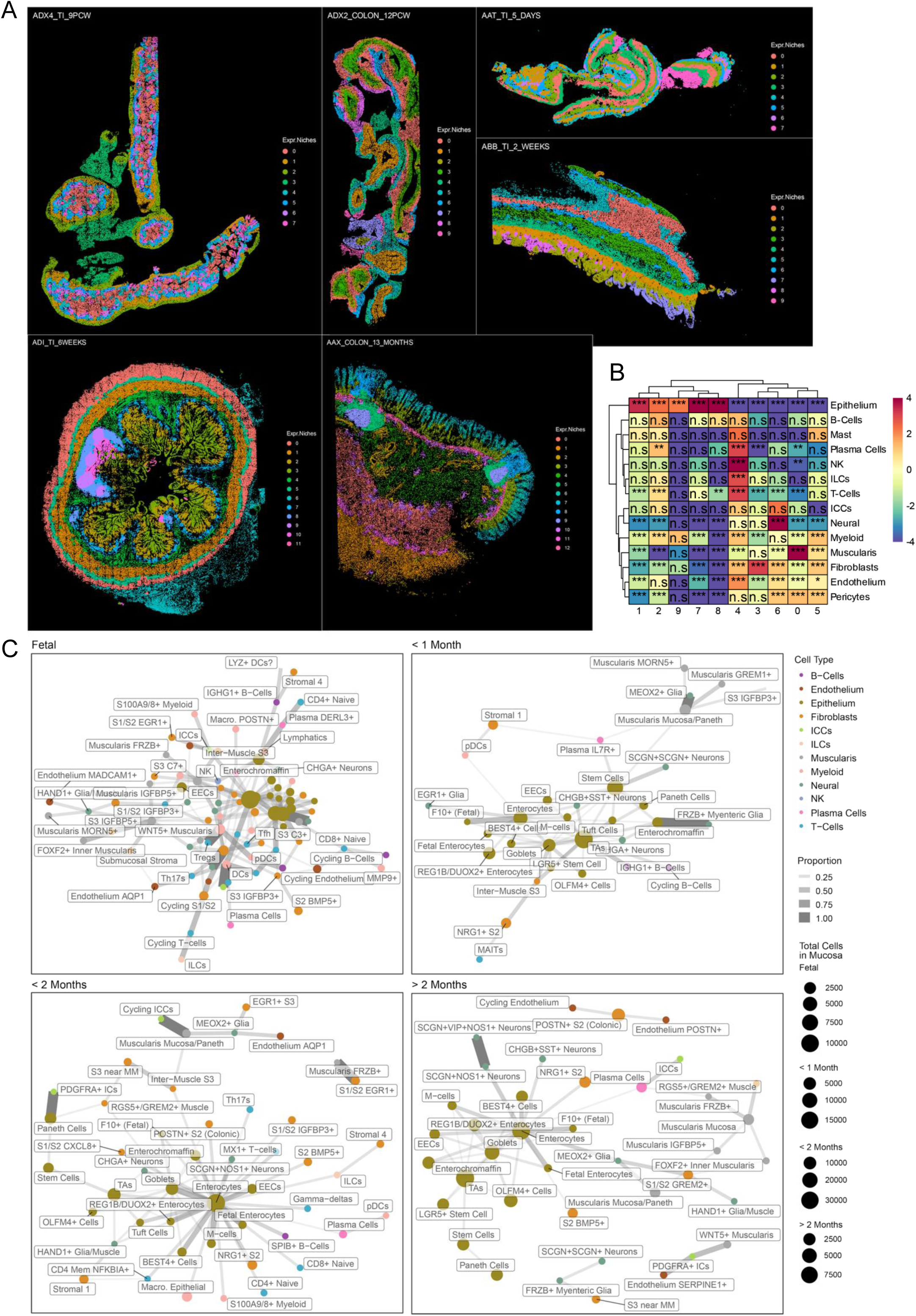
Spatial niche organisation and cell-type co-localisation. A. Representative MERSCOPE ST sections coloured by inferred spatial niches. TI-terminal ileum. B. Heatmap of cell type–niche enrichment for a representative terminal ileum sample (ABB, 2 weeks). Values are shown as log2-transformed estimates, with statistical significance indicated by asterisks (FDR < 0.05, ** < 0.01, *** < 0.001; n.s., not significant). C. Cluster co-localisation networks within the crypt niche across developmental stages. Networks include clusters with total interaction proportion ≥ 0.2 and edges with weight ≥ 0.1. Node size reflects cell abundance, edge thickness indicates co-localisation strength, and node colour denotes cell compartment.

**Supplementary Figure 10.**
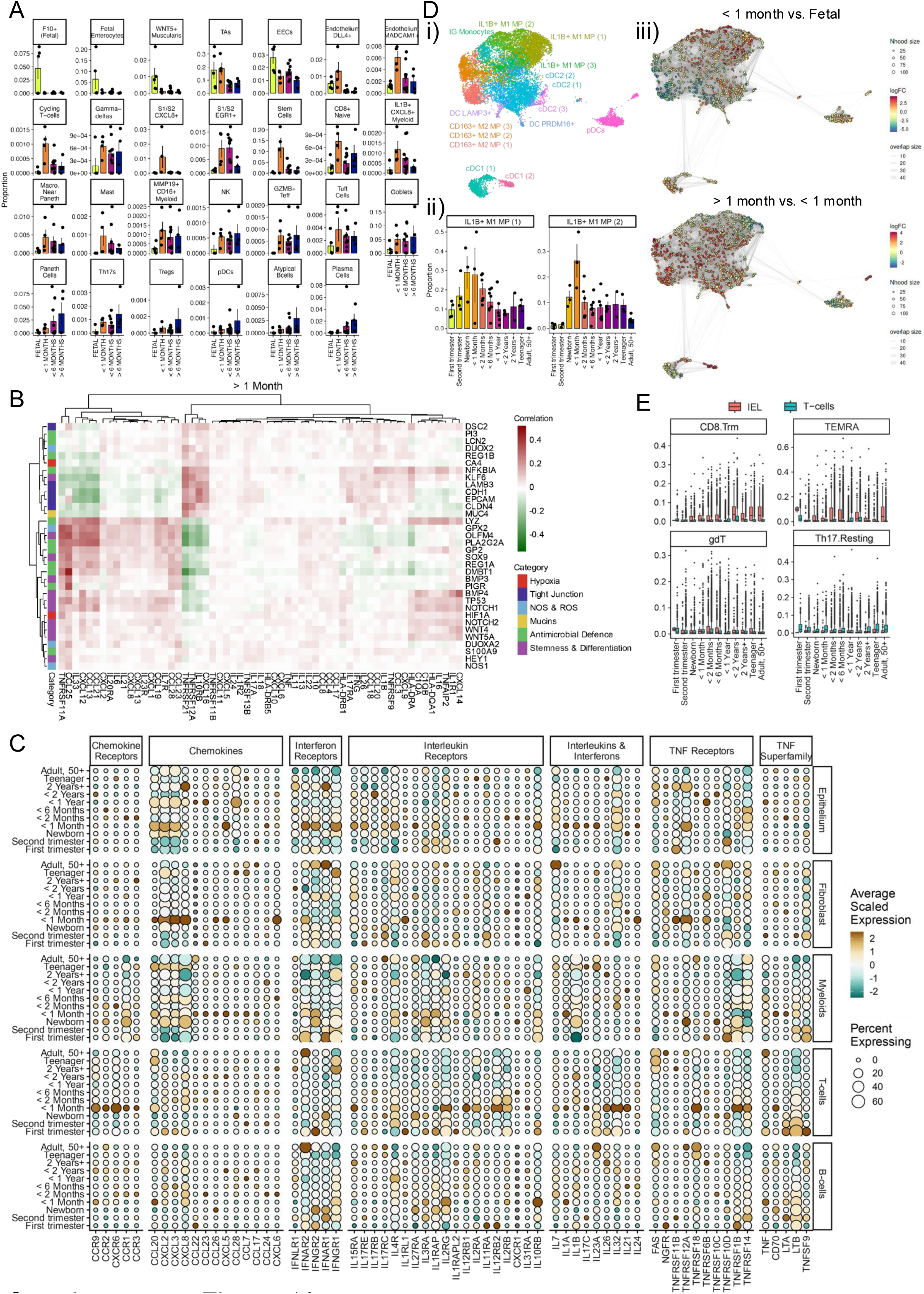
Immune composition and epithelial–neighbour interactions. A. Proportion of selected clusters in MERSCOPE ST data across age groups; points represent individual donors and bars indicate mean proportions. B. Heatmaps of epithelial–neighbourhood correlations across developmental stages. Pearson correlations between epithelial gene expression programmes and neighbouring cytokine-related signals were computed in terminal ileum samples, averaged within age groups (>1 month), and visualized by hierarchical clustering using the same colour scale as Fig. 4D. C. Bubble plot showing expression of cytokines and receptors across broad cell types in each age group. D. Age-associated changes in the myeloid compartment. (i) UMAP of myeloid cells coloured by cluster identity. (ii) Proportion of IL1B⁺ M1 macrophages within the myeloid compartment. (iii) Differential cluster abundance comparing <1 month versus fetal (top) and >1 month versus <1 month (bottom), assessed using MiloR. E. Box plots showing the distribution of T cell states, inferred using StarCAT, in intraepithelial lymphocytes from crypt-chelated epithelium and in CD45⁺ T cells. CD8 Trm, CD8⁺ tissue-resident memory T cells; TEMRA, terminally differentiated effector memory T cells; gdT, γδ T cells.

**Supplementary Figure 11.**
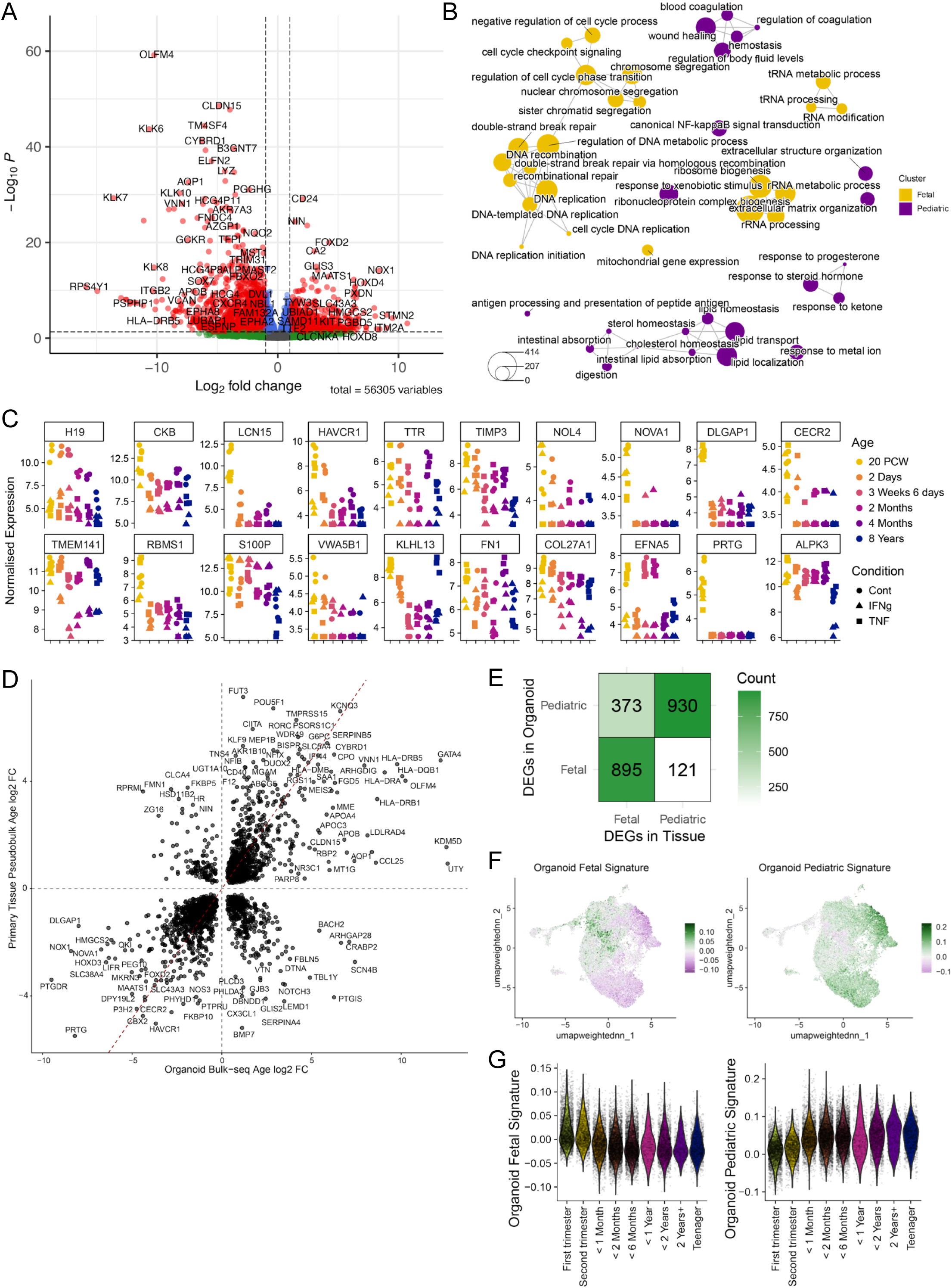
Comparison of fetal and paediatric transcriptional programmes in organoids and primary tissue. A. Volcano plot showing differential expression between fetal and paediatric organoids under unstimulated conditions, assessed using DESeq2 (Wald test with Benjamini–Hochberg correction). Points are coloured by significance and effect size. B. Network representation of pathways enriched among fetal and paediatric differentially expressed genes. C. Scatterplot showing normalized expression of fetal marker genes in organoids, coloured by age and shaped by stimulation condition. D. Comparison of fetal versus paediatric differential expression between ileal epithelial primary tissue (pseudobulk multiomics) and organoid bulk RNA-seq data. Scatterplot shows log2 fold changes; the dashed line indicates x = y. E. Grid showing the number of differentially expressed genes (DEGs) identified in fetal and paediatric samples in primary tissue and organoids. F. UMAP projection of organoid-derived fetal and paediatric gene signatures onto primary ileal epithelial multiomics data. G. Violin plots showing organoid-derived fetal and paediatric signature scores across age groups in primary ileal epithelial multiomics data.

**Supplementary Figure 12.**
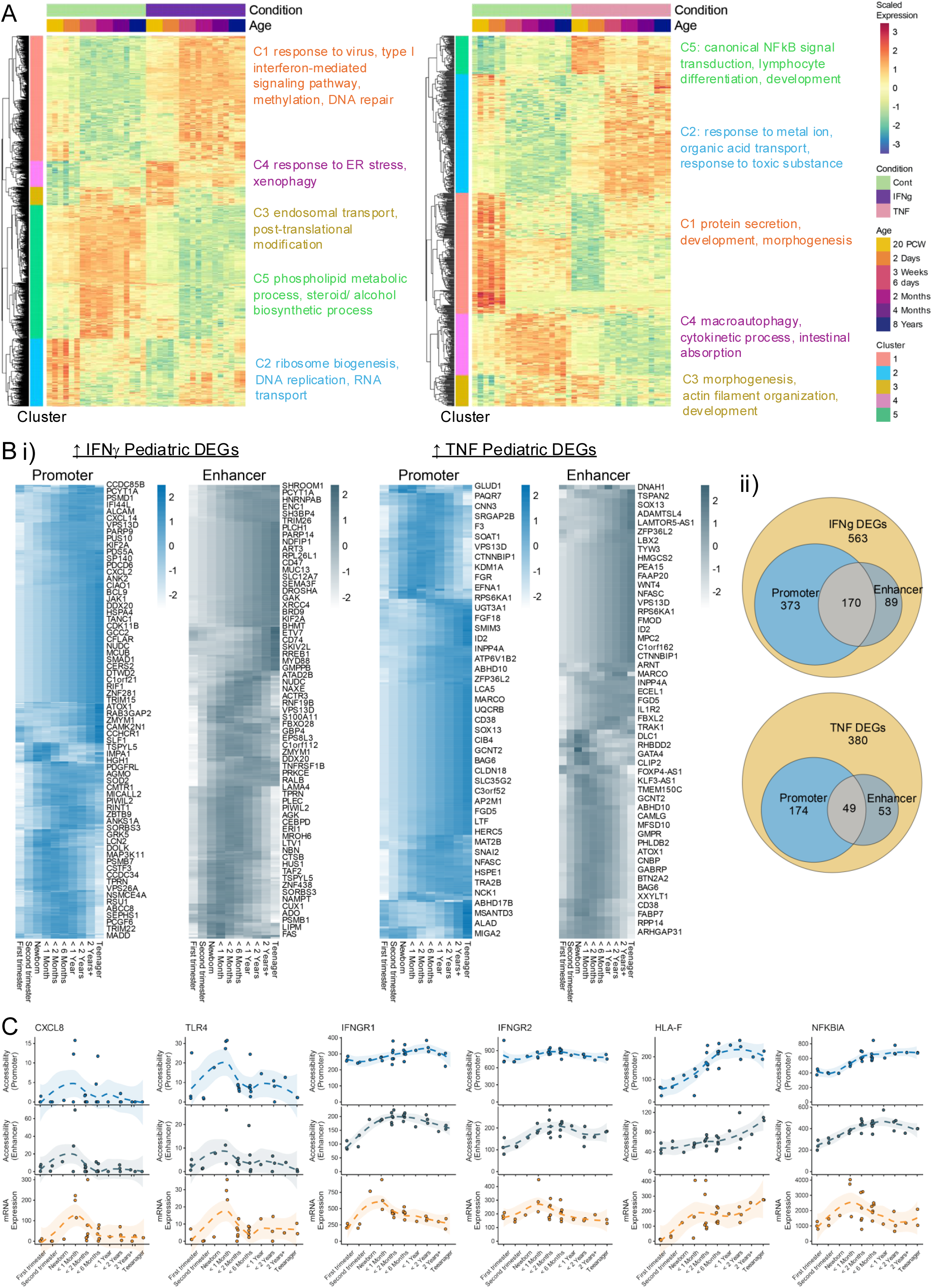
Cytokine-induced transcriptional changes in organoids and their accessibility in primary tissue. A. Heatmaps showing scaled average expression of genes significantly altered by cytokine stimulation or stimulation × age interaction (IFNγ, left; TNF, right). Genes are clustered by expression pattern and annotated with enriched Gene Ontology (GO) pathways. B. Comparison of gene expression and chromatin accessibility changes in ileal epithelial single cell multiomics data for cytokine-responsive genes identified in paediatric organoids. (i) Heatmaps showing mean GAM-fitted promoter and enhancer accessibility across age groups. (ii) Venn diagrams showing overlap of genes with increased expression in paediatric samples following cytokine stimulation, stratified by promoter and enhancer accessibility. C. GAM-fitted trends of promoter and enhancer accessibility and gene expression per sample for selected genes across age in ileal epithelium. Shaded areas indicate +/- 1 standard error of the GAM fitted mean.

**Supplementary Figure 13.**
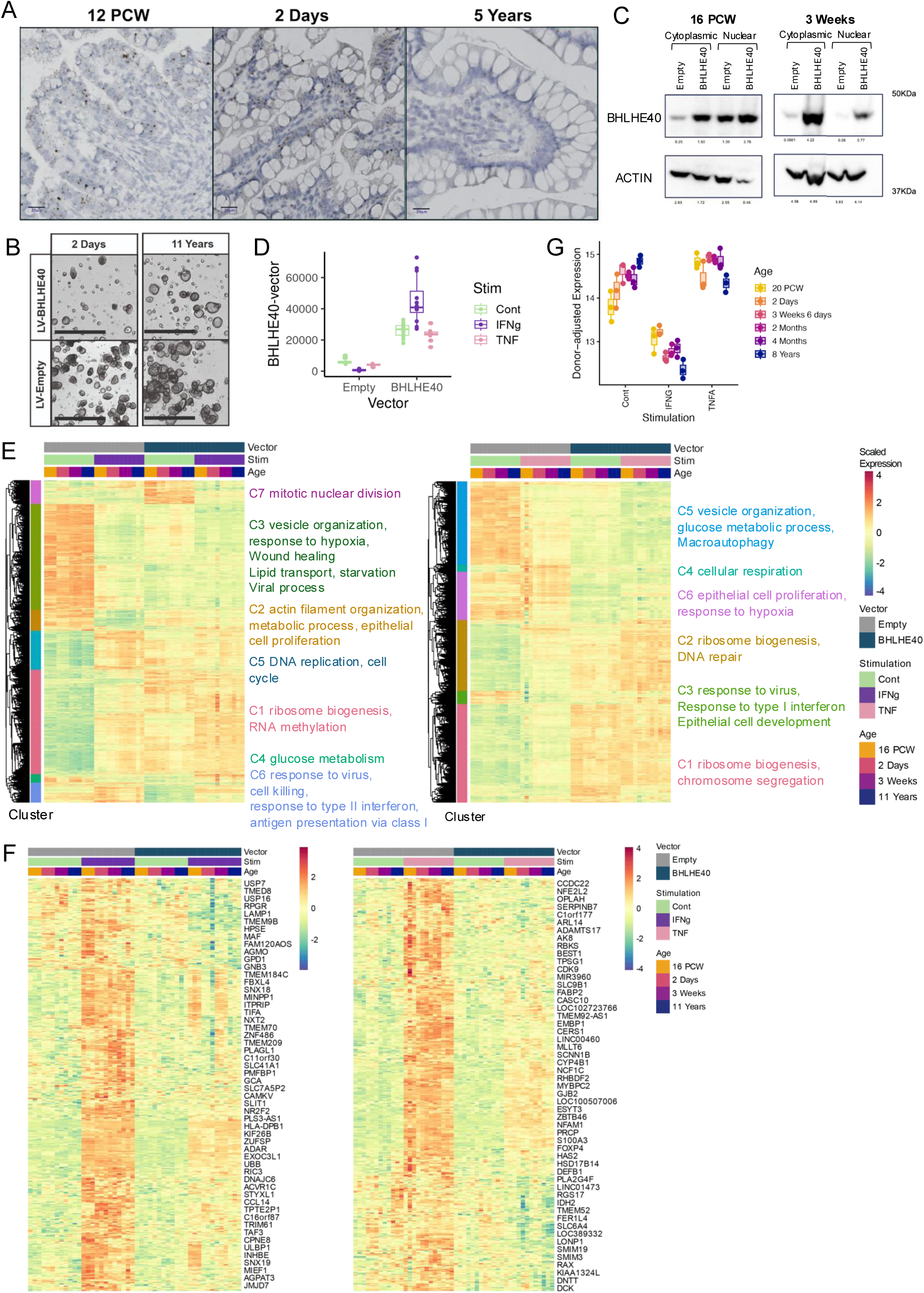
BHLHE40 expression and functional effects in organoids. A. Representative in situ hybridization (ISH) images showing BHLHE40 expression across developmental stages. Scale bars, 20 μm. B. Representative images of organoids transduced with control or BHLHE40-expressing vectors. Scale bars, 250 μm. C. Validation of BHLHE40 overexpression by western blot. D. Expression of the BHLHE40 vector in bulk RNA-seq data. E. Heatmaps showing scaled average expression of genes significantly altered by BHLHE40 overexpression and/or cytokine stimulation (IFNγ, left; TNF, right), clustered by expression pattern and annotated with enriched GO pathways. F. Heatmap showing scaled average expression of genes selectively upregulated in the empty vector condition upon stimulation (IFNγ, left; TNF, right) and attenuated with BHLHE40 overexpression. G. Scatterplot showing donor-adjusted BHLHE40 expression in organoid stimulation experiments, coloured by age.

